# Building A Unified Model for Drug Synergy Analysis Powered by Large Language Models

**DOI:** 10.1101/2024.04.08.588634

**Authors:** Tianyu Liu, Tinyi Chu, Xiao Luo, Hongyu Zhao

## Abstract

Drug synergy prediction is a challenging and important task in the treatment of complex diseases including cancer. In this manuscript, we present a novel unified Model, known as BAITSAO, for tasks related to drug synergy prediction with a unified pipeline to handle different datasets. We construct the training datasets for BAITSAO based on the context-enriched embeddings from Large Language Models for the initial representation of drugs and cell lines. After demonstrating the relevance of these embeddings, we pre-train BAITSAO with a large-scale drug synergy database under a multi-task learning framework with rigorous selections of tasks. We demonstrate the superiority of the model architecture and the pre-trained strategies of BAITSAO over other methods through comprehensive benchmark analysis. Moreover, we investigate the sensitivity of BAITSAO and illustrate its unique functions including new drug discoveries, drug combinations-gene interaction, and multi-drug synergy predictions.

## 1 Introduction

Treating patients with a combination of drugs has become common for various diseases, including HIV [1] and cancers [2, 3]. One key aspect of drug combinations is the synergistic effect, which means that the joint effect of multiple drugs is larger than the sum of individual drug effects [4]. Other definitions have also been used to define synergistic effects, such as [5]. Effective drug combination can reduce drug resistance of monotherapy [6] with relatively lower doses of individual drugs [7]. Since drugs can change gene expressions when applied to different systems, e.g., cell lines, their effects can be studied through the genomics lens [8, 9]. Currently, researchers use high-throughput combinatorial screening to identify drug combinations with synergistic effects for specific cell lines [10]. However, such experimental screening is laborious and time-consuming due to the very number of potential drug combinations, and it is even more challenging to assess the synergistic effect of combinations with three or more drugs [11]. Therefore, it is important to develop computational methods based on extensive experimental datasets in the public domain as well as diverse types of prior biological knowledge to predict the presence and strength of synergistic effects for candidate drug combinations. Accurate prediction methods can facilitate drug discovery [12] and clinical development [13].

Given its importance, it is no surprise that many machine learning methods, especially deep learning methods, have been proposed to predict drug synergy. These methods differ in model architecture, training strategies, and datasets used to build the models. DeepSynergy [14] is among the earliest tools by building a neural network for both regression and classification, with follow-up work such as TreeComb [15, 16] and MatchMarker [17]. Existing drug synergy prediction methods can be broadly classified into two groups. The first group of methods, such as MARSY [18], focus on predicting specific synergy scores, whereas the second group of methods, such as DeepDDs [19], transfer the continuous synergy score into a binary one via thresholding to infer drug combination synergy. However, most existing methods do not incorporate the extensive synergy information from public databases [20] in their predictions. [21] utilized a transfer learning approach and pre-trained the model based on large-scale databases while incorporating different types of features (e.g., gene expression, molecular structure). However, it did not consider datasets [14] with only partial information and treated drugs with the same molecular formula but different names as distinct ones. Therefore, the generalization ability of this model is limited by its input data format. Moreover, since public databases are updated constantly, it is important to track the versions of training datasets.

Large Language Models, as a type of Foundation Models (FMs) [22], have greatly improved the performance of deep learning on various tasks in Natural Language Processing (NLP) [23]. Such models have received broad attention from both industry and academia [24]. Researchers have proposed to use FMs to predict drug synergy via LLMs by transferring the drug synergy prediction problem into a Question-Answer problem [25, 26]. By incorporating prior information of single drug and single cell line from LLMs, it has become possible to predict drug synergy of unknown drug combinations in unknown cell lines. Text information may be less noisy than the features (e.g. gene expression levels) that have been used in this task. However, such QA setting limits the task to a classification problem, which introduces the potential bias of pre-defined thresholds. Moreover, these two LLMs are not open-source so it is difficult for researchers to evaluate their performances. Open-source is important for the development of science [27]. Moreover, there is a lack of exploration on the utilization of the information in LLMs for more difficult drug synergy prediction problems, e.g., the effects of multiple drug combinations or model explainability.

Here we present a scalable unified model for drug synergy prediction called BAITSAO^1^. BAITSAO utilizes the information from LLMs as input and was pre-trained based on large-scale known synergistic effect information of paired drug combinations and cell lines. The information of drug combinations and cell lines is necessary to predict synergy scores. We show that the embeddings of these features from LLMs can be effective input for drug synergy prediction as well as the effects of drugs on gene expressions. We further demonstrate the capability of building an effective predictor for synergy prediction under both the classification and regression settings through multi-task learning (MTL) [28]. Finally, we pre-train BAITSAO to predict synergistic effects for unseen drug combinations based on the zero-shot learning framework and the fine-tuning framework. The scalability of BAITSAO allows us to consider multiple drugs and incorporate extra meta information.

## 2 Results

### Overview of BAITSAO

We highlight two major contributions of BAITSAO as a unified model. We first provide a new unified pipeline for pre-processing the information from both drugs and cell lines for machine learning in a tabular format, and generate training datasets from these embeddings for multiple tasks. We show that these embeddings contain functional information for prediction. We then utilize the unified training datasets for different synergistic effect prediction tasks under the multi-task learning framework. We demonstrate the superiority of the model architecture and the contribution of pre-training through comprehensive experiments. BAITSAO can be easily transferred to perform novel downstream tasks related to drug synergy analysis. We illustrate the landscape of BAITSAO in Figures 1 (a) and (b) and summarize the differences between BAITSAO and other synergy prediction methods in Figure 1 (c). The major functions of BAITSAO are shown in Figure 1 (d).

**Fig. 1.**
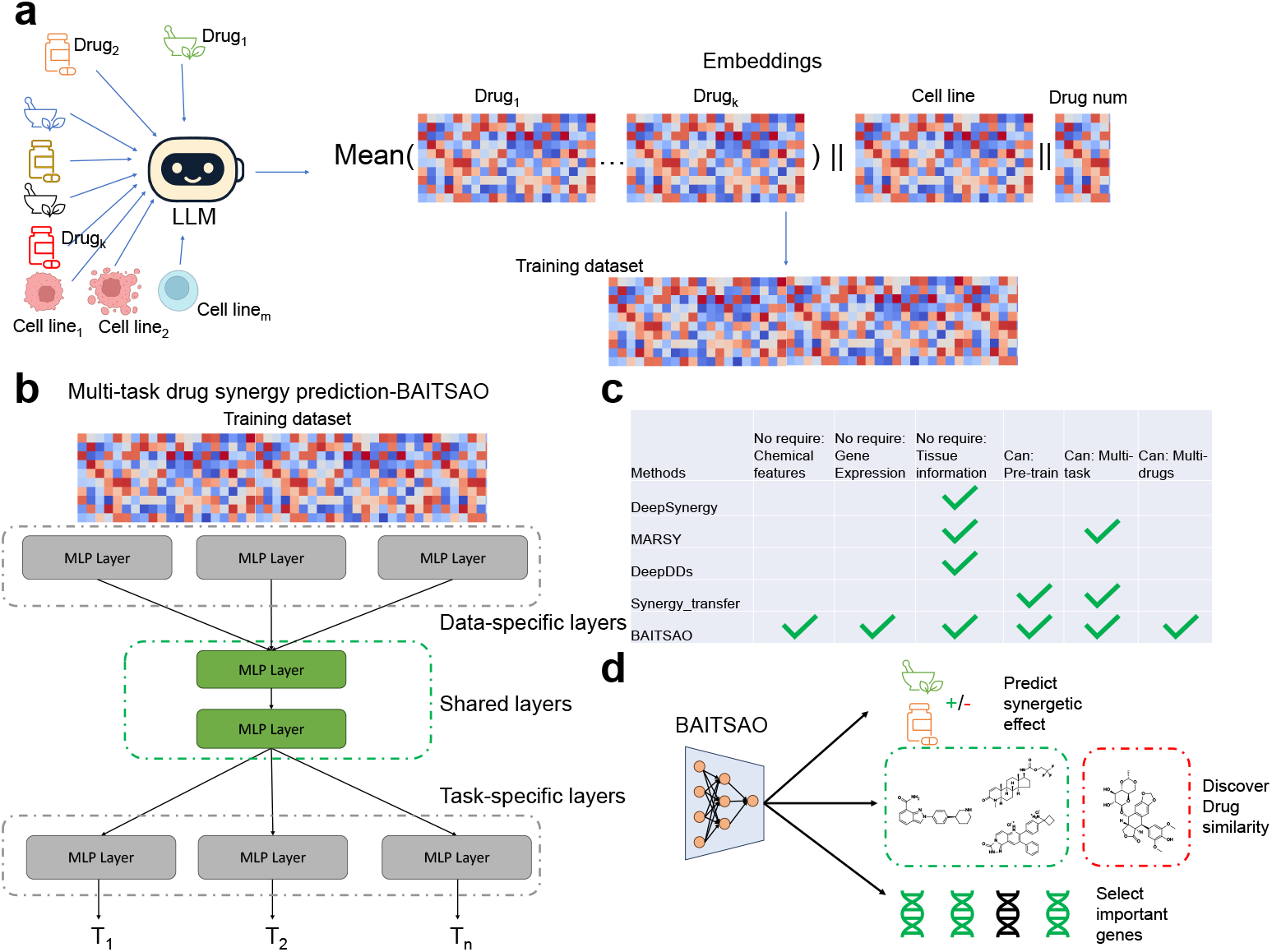
An overview of BAITSAO as a FM under the pre-training and fine-tuning/zero-shot learning pipeline. (a) The pre-processing steps we used to transfer the meta information into embeddings to construct training datasets. (b) The model architecture of BAITSAO under a multi-task learning framework. (c) Comparisons of different methods for drug synergy analysis. (d) Different functions of BAITSAO.

### Drug embeddings from LLMs reflect functional similarity and responses at the cell level

In this section, we discuss the information offered by drug embeddings and cell-line embeddings. We generate the description for the drugs and cell lines from our training datasets based on designed prompts from LLMs, and then use the embedding module from GPT 3.5 [29] to generate the embeddings of such descriptions, where the embeddings become the features of drugs or cell lines. We utilized GPT 3.5 rather than GPT 4 [30] because the layer for generating embeddings is from GPT 3 [29] series and the querying time from GPT 4 with similar quality required much more time [31], and efficiency is very important in LLM deployment [32, 33]. Moreover, the performance difference between embeddings from GPT 3.5 and GPT 4 or from GPT 3.5 and Claude 3.5 [34] is not significant based on our experiments, shown in Supplementary Figure 1 (a) (Wilcoxon rank-sum test, p-value=0.86 for GPT 3.5 vs. GPT 4, and p-value=0.44 for GPT 3.5 vs. Claude 3.5). In the same figure, we also found that embeddings from GPT 3.5 are better than embeddings from Gemini [35] (p-value=0.0039), and thus our current selection is well-designed. We visualize the drug embeddings and the cell-line embeddings based on Uniform Manifold Approximation and Projection (UMAP) [36] shown in Supplementary Figures 2 (a) and (b). We investigated the quality of the embeddings by considering both the quality of the description and the quality of the functions of embeddings.

For the first aspect, we recorded the outputs as descriptions from GPT 3.5 based on our prompts and compared the content with information from DrugBank [37] and NCBI [38]. Here we used drugs and cell lines from DeepSynergy, which contains 39 drugs and 38 cell lines. The descriptions summarized the functional information of drugs and cell lines. Based on our experiments, only one drug (MK-8669) has a mis-matched generated description, while 13 drugs cannot be matched with indication information if we search them in DrugBank. All descriptions are included in Supplementary file 1. We plot the Cosine Similarity (CS) for all drugs’ embeddings in Figure 2 (a). We also randomly selected 10 drugs from this dataset and plot the CS for the embeddings of the same drug under 10 different descriptions by running GPT 3.5 multiple times in Supplementary Figure 3. These two figures show that the similarity from different drugs is generally lower than that from the same drug, suggesting that we can get informative embeddings from LLMs.

**Fig. 2.**
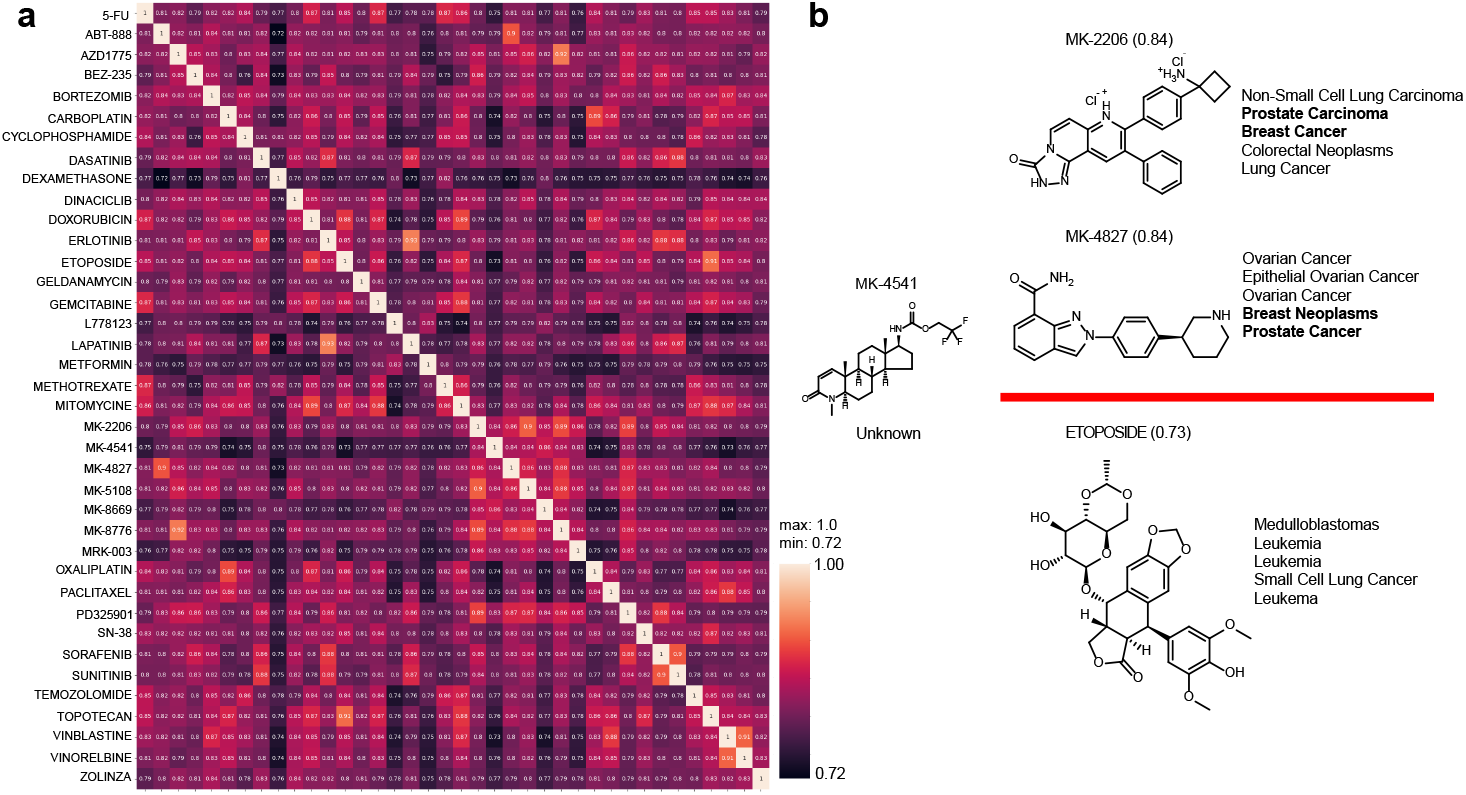
Investigation of drug embeddings. (a) The heatmap for the similarity of embeddings across all the drugs. (b) Exploration of drug similarity related to MK-4541. The drugs above the red line represent the two most similar drugs, while the drugs below the red line represent the most different drug. We list five types of clinical trial information ranked by the phases. Source data are provided as a Source Data file.

To perform a comprehensive analysis of our generated drug embeddings from LLMs, we downloaded the descriptions of drugs, including indication, summary, and background, from the DrugBank. We embedded these descriptions based on the same GPT-3.5 embeddings layer and computed the CS between embeddings from Drug-Bank descriptions and the LLM-generated descriptions. We found that embeddings from LLMs have a strong average similarity with all three descriptions from DrugBank (CS=0.87 for indication, CS=0.90 for summary, and CS=0.90 for background), and thus the generated drug embeddings preserved the important functional and chemical properties of the original drug. Furthermore, we visualize the CS based on the embeddings from drug indication (Supplementary Figure 4 (a)), drug summary (Supplementary Figure 4 (b)) and drug background (Supplementary Figure 4 (c)). We further computed the Pearson Correlation Coefficient (PCC) between the similarity matrix from DrugBank descriptions and LLM descriptions, which could be used to evaluate the ability of embeddings used by BAITSAO in preserving the drug-drug similarity. The PCCs are annotated under each figure, and all of the PCCs are high (PCC ≥ 0.76) and significant (p-value*<*0.05). Therefore, we demonstrated the ability of LLMs to generate meaningful descriptions as well as embeddings by comparing the generated information with known database, and further enhanced the reliability of the pipeline.

Furthermore, we performed Mann-Whitney U test [39] to compare the PCCs among the drugs from the MK class and the PCCs between the drugs from the MK class and other classes, and the test statistics showed a significant difference (p-value=9.9e-12). Therefore, in Figure 2 (b), we used drug MK-4541 as one example and there is no clinical information for this drug in the DrugBank, to infer its function based on our embeddings. By excluding the drug MK-8669 due to mismatched information, drug MK-2206, and drug MK-4827 have the highest similarity with MK-4541. Since MK-2206 and MK-4827 have similar functions (e.g., treating Breast-cancer-related and Prostate-cancer-related diseases), we may infer that MK-4541 may have a similar effect. Among these drugs, EPTOPOSIDE has the lowest similarity and it also has different clinical trial information, suggesting that correlation between embedding similarity and function similarity. Therefore, our drug embeddings may help the inference of clinical functions of drugs based on the embeddings’ similarity.

To investigate whether the embeddings can be used to predict drug response for the cell-level task, we utilized CPA [9] and single-cell RNA sequencing (scRNA-seq) [40] datasets with different perturbations (defined by different drugs or drug combinations) to evaluate whether our drug embeddings can facilitate the gene expression prediction task. With drug embeddings, we can use CPA to predict gene expression response to unseen drugs. Cells with unknown perturbation results are also known as out-of-distribution (OOD) samples. The original implementation of CPA utilized the drug embeddings from Rdkit [41, 42] to encode the molecular structure of the selected drug into the embedding space. However, such methods could not handle drugs not in the Simplified Molecular-input Line-entry System (SMILES) [43], which limits the generalization of CPA. Here we considered replacing the original embeddings in CPA with the embeddings from GPT 3.5, enlarging the accessibility for drug embeddings. We compared three different embedding settings for two datasets (CPA example [9] and Openproblems [44]), which contain the gene expression profiles under the control case and drug-based perturbations. The results are shown in Supplementary Figures 5 (a)-(d), where stacking the embeddings from SMILES and GPT 3.5 achieved the best performance under both datasets. For the CPA dataset, both using the embeddings from GPT 3.5 and the setting of embeddings stacking can enhance the prediction performance significantly, compared with the mode of only using SMILES (Wilcoxon rank-sum test, p-values*<*0.05). For the Openproblems dataset, the contribution of such embeddings stacking for prediction is especially significant (p-values*<*0.05). Therefore, the embeddings from LLMs can improve the gene expression prediction for perturbed scRNA-seq data.

Since our experiments demonstrate that drug embeddings and cell embeddings can summarize the functional information and drug embeddings can also interact with cell-level gene expressions, we believe that these embeddings allow us to construct the training dataset to predict drug synergy effect in different cell lines.

### Demonstration of powerful embeddings and architecture by evaluation without pre-training

In this section, we show the strength of LLM embeddings and select the choice of network structure for model pre-training based on two different drug synergy prediction tasks: classification and regression. For each task, we selected two datasets and two metrics for evaluation. For regression, we included the Pearson Correlation Coefficient (PCC) and Mean Squared Error (MSE) for model evaluation based on datasets D1 [14] and D2 [18]. For classification, we included ROC-AUC (ROCAUC) and Accuracy (ACC) for model evaluation based on datasets D1 and D3 [19]. These metrics and datasets were widely used in the related work [14, 18, 19, 45, 46]. First, we tested if the model performance will be affected by prompt engineering of LLMs, and we compared the raw embeddings with embeddings generated by drug descriptions from MetaPrompt [47] and Chain-of-Thought (COT) [48]. According to Supplementary Figure 1 (b), the differences between the default mode and these two prompt engineering methods are not significant (p-value=0.63 for raw mode vs. MetaPrompt, and p-value=0.43 for raw mode vs. COT). Therefore, our embeddings have enough information as inputs for synergetic effects. Second, we validated the contribution of BAITSAO’s architecture shown in Supplementary Figure 1 (c). We compared the performances between BAITSAO and DeepSynergy with LLM embeddings as inputs. The difference is significant and thus our optimization of model architecture also contributed to the prediction task (p-value=0.002). Finally, we selected seven other methods (DeepSynergy, MARSY, TreeComb, SVM [39, 49], TabNet [50], BERT [51] and Lasso [39, 52]) for benchmarking the regression task and seven methods (DeepSynergy, DeepDDs, TreeComb, SVC, TabNet, BERT, and Lasso) for benchmarking the classification task. We utilized the best hyper-parameters of these methods for every dataset, with details of hyper-parameter tuning summarized in the Methods section. Our results based on five-fold cross-validation [14] are summarized in Figure 3. This figure shows that BAITSAO ranked the best in three out of four metrics. Moreover, BAITSAO was also the most stable among the top deep-learning-based methods (including MARSY, DeepSynergy, DeepDDs, and TabNet). The performance of BERT was worse than BAITSAO in three out of four metrics, thus using embeddings as input is better than using the combination of description in general. For the evaluation based on MSE, BAITSAO performed well on the D1 dataset. Our experiments showed that embeddings from LLMs with a suitable model architecture can formalize a better training-testing framework compared with data from the classical feature space. The details of our dataset information, model construction, and training process are summarized in the Methods section.

**Fig. 3.**
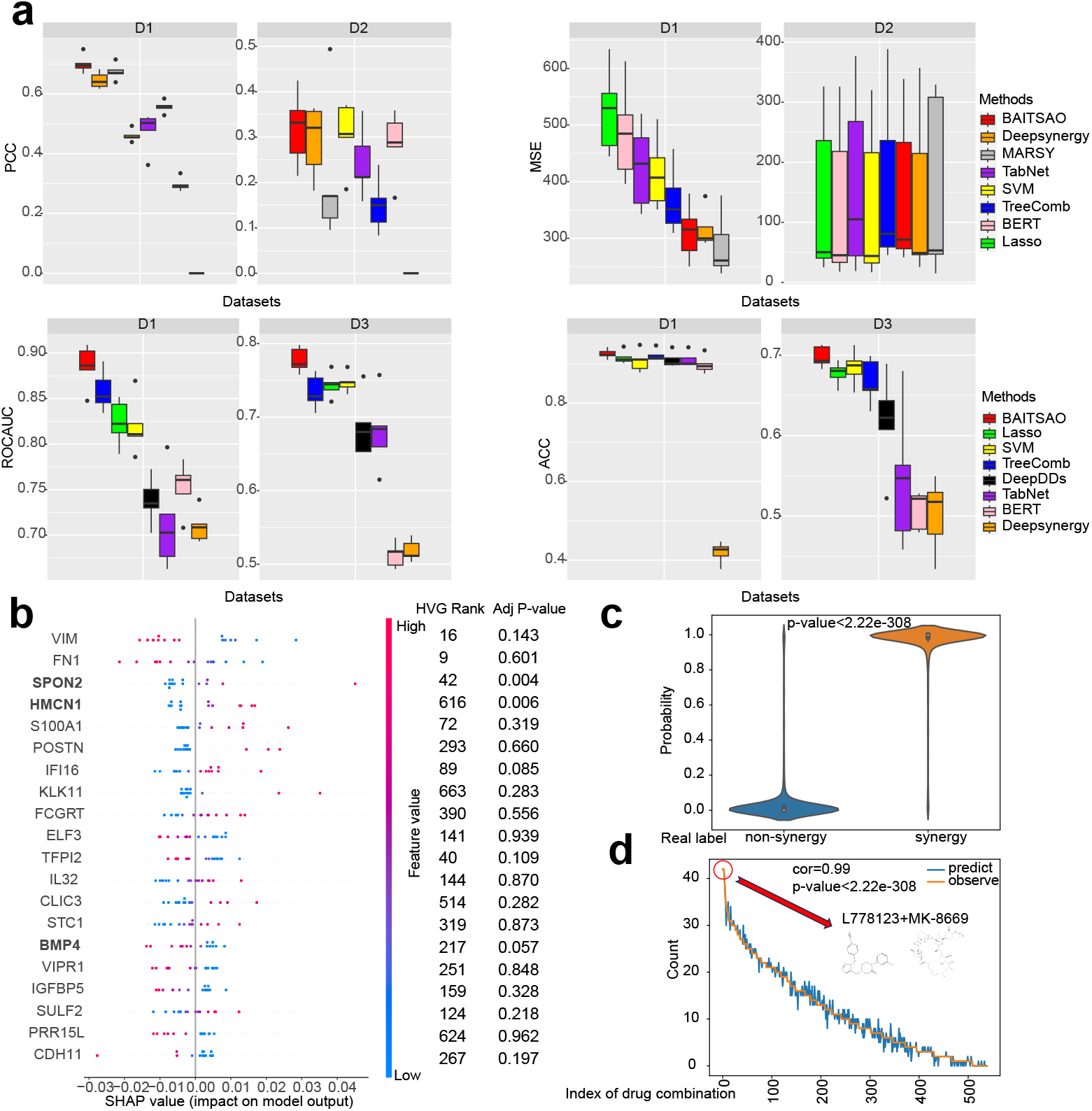
Results of evaluations for model structure, reliability and explainability. (a) Evaluations for BAITSAO and other methods. Each panel represents one metric with two datasets. The ranks of methods are averaged by datasets. Data are presented as boxplots (n=5 per group; center line, median; box limits, upper and lower quartiles; whiskers, up to 1.5 *×* interquartile range; points, outliers). The explanations of datasets D1-D3 are summarized in the Methods section. (b) The explainability of BAITSAO for the combination DEXAMETHASONE (drug)-DINACICLIB (drug) for different cell lines. We also list the ranks based on variance (HVG rank) and the adjusted p-value based on DESeq2 analysis results for each gene. The genes with adjusted p-value for multiple comparisons smaller or close to 0.05 are boldfaced. (c) The violin plot (n=6299 for non-synergy group; n=6116 for synergy group; center point, median; box limits, upper and lower quartiles; whiskers, up to 1.5 *×* interquartile range; points, outliers) for the outputs of BAITSAO (Probability) across the synergistic labels. We also present the two-side p-value in this figure. This panel supports the reliability of selected features from SHAP. (d) The rank-based plot between the number of drugs-cell line combinations with synergy (Count) and the index of drug combination (Index of drug combination). The index is ranked by the value of Count. We present the Pearson correlation (corr) and two-side p-value in this figure. Source data are provided as a Source Data file.

### Explainability of BAITSAO for drug-gene interaction and drug-cell line interaction with multi-modal learning

We interpret contributions of different features for the prediction task with the help of SHAP [53]. Here we integrated known gene expression profiles of different cell lines in D3 to our input datasets and performed the same training process for the drug synergy prediction task. We then utilized SHAP to study the importance of different genes and the results could be treated as the relevance between the gene expression levels (as a new modality) and the possibility to produce synergistic effect for drug combinations. We followed the default setting of SHAP to fix the number of genes for explainability at 20. We also performed statistical analysis based on the outputs of BAITSAO to discover the drug combination with the largest range of synergistic targets. The details of our approach are provided in the Methods section.

By collecting gene expression profiles of cell lines [54], we studied the explainability of BAITSAO for DEXAMETHASONE (drug)-DINACICLIB (drug) across different cell lines. In Figure 3 (b), we visualize the importance of different genes. The gene VIM was top-ranked by the average importance, and VIM is known as important for various cancers from pan-cancer analysis [55]. Furthermore, we conducted three experiments to further investigate the contributions of the selected genes.

We first separated the samples into two groups based on the existence of the synergistic effect and performed DEG analysis using DESeq2 [56, 57] between two groups of cell lines. We present the adjusted p-values using Benjamini-Hochberg for the selected genes in Figure 3 (b). Genes SPON2, HMCN1, and BMP4 listed in this figure were significant DEGs. Our selected genes had significant overlap with DEGs (Fisher’s exact test p-value=0.0062), and the gene BMP4 is a validated targets for each drug according to biology experiments [58, 59]. Moreover, these genes had relatively lower expression levels with synergistic effect, which matched the distribution of their SHAP values (enriched in the negative values).

We list the ranks of the selected genes based on their variances in Figure 3 (b). The top five ranked genes had relatively greater variances, suggesting that the genes we selected characterize the heterogeneity from both cell lines and the drug synergistic effect. We further performed enrichment analysis based on Gene Ontology (GO) [60] for biological pathways and Molecular Signature Database (MsigDB) [61] for cancer-specific signals based on this set of genes, with results shown in Supplementary Figures 6 (a) and (c). These enriched pathways represent important biological processes and cancer-specific signals. These results suggest that our method may uncover the heterogeneity in the drug synergy prediction process. We summarize our results for the single cell line with the same drug combination in Appendix A. The plots for important genes across different cell lines can be found in Supplementary Figure 16. The test statistics used in this section are given in Supplementary file 2.

We further investigated the drug combination that showed synergistic effects on the largest number of cell lines. We first plot the probabilities of all drug-cell line combinations to be classified as samples with synergistic effects in Figure 3 (c). This figure shows that the distribution of such probability is different under different synergistic labels. We performed the Rank-sum test [62] for these two sets of probabilities and their difference is significant (p-value*<*2.22e-308 with two-side mode and no adjustment is needed). Therefore, our model can uncover the relationship between input features and the synergistic effect. Moreover, we ranked the drug combinations based on the number of cell lines predicted to have synergistic effects in descending order. We computed the Pearson correlation coefficient [62] between the count value based on predicted labels and observed labels, summarized in Figure 3 (d). Based on this figure, the count values based on the predicted labels had a strong positive correlation with those based on the real labels, thus our model can also be used to identify the drug combinations with the most synergistic targets given a set of cell lines and cancer types used in our experiments. We also highlight the drug combination L778123 and MK-8669 that has the largest number of targeted cell lines with synergistic effect in Figure 3 (d). The p-value is computed with two-side mode and no adjustment is needed. Therefore, BAITSAO can capture the variance of different drug combinations across cell lines, offering a novel option for selecting effective drug combinations.

### Statistics of pre-training datasets

Here we summarize the statistics and properties of our pre-training datasets for BAITSAO. We collected information from DrugComb [20], which is known as the largest database containing synergistic effect information for drug pairs with different cell lines. We downloaded the updated version of DrugComb and removed the missing value or single-drug information. The major statistics of DrugComb are summarized in Figure 4, whose (a) represents the total number of *drug-cell line* combinations by tissue types and Figure 4 (b) represents the total number of *cell lines* by the type of tissues from DrugComb. Most of the drugs presented here were analyzed using cells from skin, lymphoid, and/or lung. These tissues are important for maintaining normal physiological activity in the human body. In total, DrugComb collects more than 700,000 available combinations. As shown in Figure 4 (c), the distribution of the synergy scores is not balanced, with a large number of combinations having low synergy scores. We further plot the Half Maximal Inhibitory Concentration (IC 50) for all drugs in Figure 4 (d) with a similar distribution to the synergy score. We illustrate the non-linear relationship between single drug IC 50 and synergy score in Figure 4 (e). Therefore, fitting non-linear models like neural networks may help the synergy prediction task. Finally, Figure 4 (f) shows the overlap of combinations by tissues, where most tissues have low overlap, and thus the pre-training dataset has information from diverse tissues. We plot the embeddings for drugs and cell lines in the pre-training dataset colored by clusters from Leiden [63] in Supplementary Figures 7 (a) and (b). The items in the same Leiden cluster can be treated in similar context of embeddings with functional information, so we can visualize the functional similarity of different drugs and cell lines through embeddings. Since our pre-training dataset was published in June 2021 and GPT 3.5 collected data for pre-training until Sep 2021, considering the time needed for pre-training a LLM, we believe that the data from DrugComb were precluded in GPT 3.5.

**Fig. 4.**
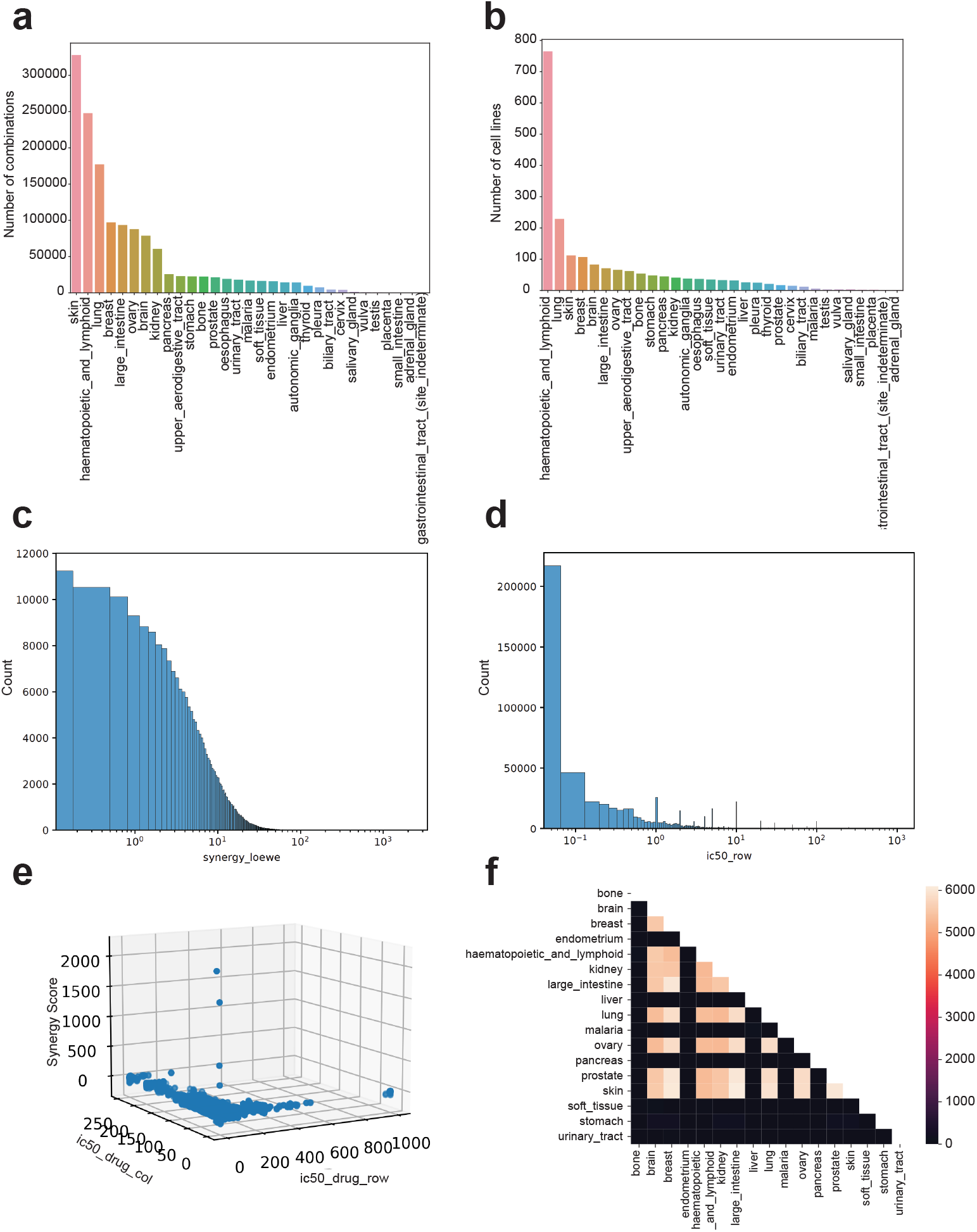
Statistics of the pre-training dataset from DrugComb. (a) The barplot for the number of *drug-cell lines* combinations by different tissues. (b) The barplot for the number of *cell lines* by different tissues. (c) The histogram for the distribution of synergy score computed based on Loewe [64]. The x-axis is transferred into log scale. (d) The histogram for the distribution of single-drug IC 50 levels. The x-axis is transferred into log scale. (e) 3D plot for the relation between single-drug IC 50 levels and synergy score. (f) The heatmap for the overlap of combinations across different tissues. Source data are provided as a Source Data file.

### Pre-trained BAITSAO contributes to drug synergistic effect prediction under the multi-task condition

Here we investigated and pre-trained BAITSAO based on the optimal model structure. Specifically, we extended the model structure with a multi-task learning framework. By pre-training BAITSAO with large-scale synergy datasets, BAITSAO is able to predict both single drug inhibition and drug synergistic effect. For drug pairs, we expect to predict both drugs’ inhibition, thus we have a total of four tasks inspired by the pre-training datasets, including the regression task for synergy prediction, the classification task for synergy prediction, and regression tasks for single-drug inhibition of each drug in the drug pairs. For the regression task of synergy prediction, we only considered predicting the synergy score under the Loewe setting because we show that the synergy scores computed based on other methods are positively correlated with the Loewe score [64] in Supplementary Figure 8, and literature [14, 25] suggests using the threshold for generating a classification task from the Loewe score. For other synergy scores including Zip score [65], HSA score [66], and Bliss score [67], we pre-trained specific models and restored the pre-training weights. Instead of using the simple average of loss functions from different tasks during the training process, we introduced the Uncertainty Weighting (UW) method [68] advocated by the performance evaluations of different multi-task learning strategies from [69] and improved the numerical stability and the validation strategy of this method.

We first determined the tasks that can help each other in the multi-task learning framework by constructing the Help-Harm matrix. We sampled 1% of the pre-training dataset and trained task-specific models as well as multi-task models with paired tasks, and constructed the Help-Harm matrix shown in Figure 5 (a). According to this figure, joint training always boosts the classification task, while joint training with the classification task can help predict the synergy scores as well as inhibition levels for a single drug. Moreover, the relative inhibition (RI) information from one of the drugs in drug pairs did not show a significant contribution to other tasks, and incorporating this information reduced performance for the classification task. Since we had RI levels for both drug pairs, we removed the information of RI col in the training process, and collected three tasks in the pre-training stage. After finishing pre-training based on the sampled and full datasets, we plot the metrics for comparing the performance between BAITSAO under the STL framework and our final MTL framework in Figure 5 (b). MTL can improve the performance of BAITSAO for solving all regression-based tasks. We show the outputs from the hidden layers of BAITSAO by ground truth synergistic labels and predicted synergistic labels in Supplementary Figures 9 (a) and (b). According to these two figures, the learned drug embeddings for drugs with no synergistic effect tended to be co-embedded. Therefore, BAITSAO with the MTL framework is reasonable and superior in drug synergy analysis. Finally, we consider the generalization ability of BAITSAO with pre-trained weights. We conducted experiments based on three datasets we used in the subsection *Selection of the model structure by evaluation without pre-training* and visualized the results in Figure 5 (c). We report the metrics based on five-fold cross-validation results. According to this figure, BAITSAO with the pre-training design after fine-tuning (BAITSAO-FT) is comparable or better for the regression and classification tasks, compared with BAITSAO without pre-training (BAITSAO-ZS). When evaluating the ZS mode, we ensured that the combinations used in the pre-training stage are not used for testing. Moreover, our fine-tuning stage used fewer epochs and we froze the shared layers during the fine-tuning process, thus our fine-tuning approach was more efficient. We note the potentials of BAITSAO under the zero-shot learning framework for solving this task. For example, BAITSAO-ZS showed a high ACC score in the evaluation based on D1. Moreover, for the metrics related to classification, BAITSAO-ZS had results higher than 0.5, and thus BAITSAO under the zero-shot learning framework was better than random guessing. We also performed Rank-sum tests [62] between the pre-training dataset and fine-tuning datasets and the results are shown in Supplementary Figure 10, which demonstrated that samples in the fine-tuning datasets satisfied the OOD cases. Finally, we compared BAITSAO with other LLM-based models, discussed in Appendix B, which shows that BAITSAO also has unique advantages. We also pre-trained other deep-learning-based synergy predictors, such as DeepSynergy, DeepDDs, and MARSY based on their designed tasks and compared the fine-tuned version of these models with BAITSAO (ft). According to Supplementary Figure 11, BAITSAO still shows better performances than other baselines with either finetuning mode or from-scratch mode. Therefore, the multi-task pre-training strategy of BAITSAO is unique and contributive, which leads to consistent improvement across different datasets. In summary, the combination of MTL and the pre-training process can improve the performance of BAITSAO on tasks related to drug synergy analysis.

**Fig. 5.**
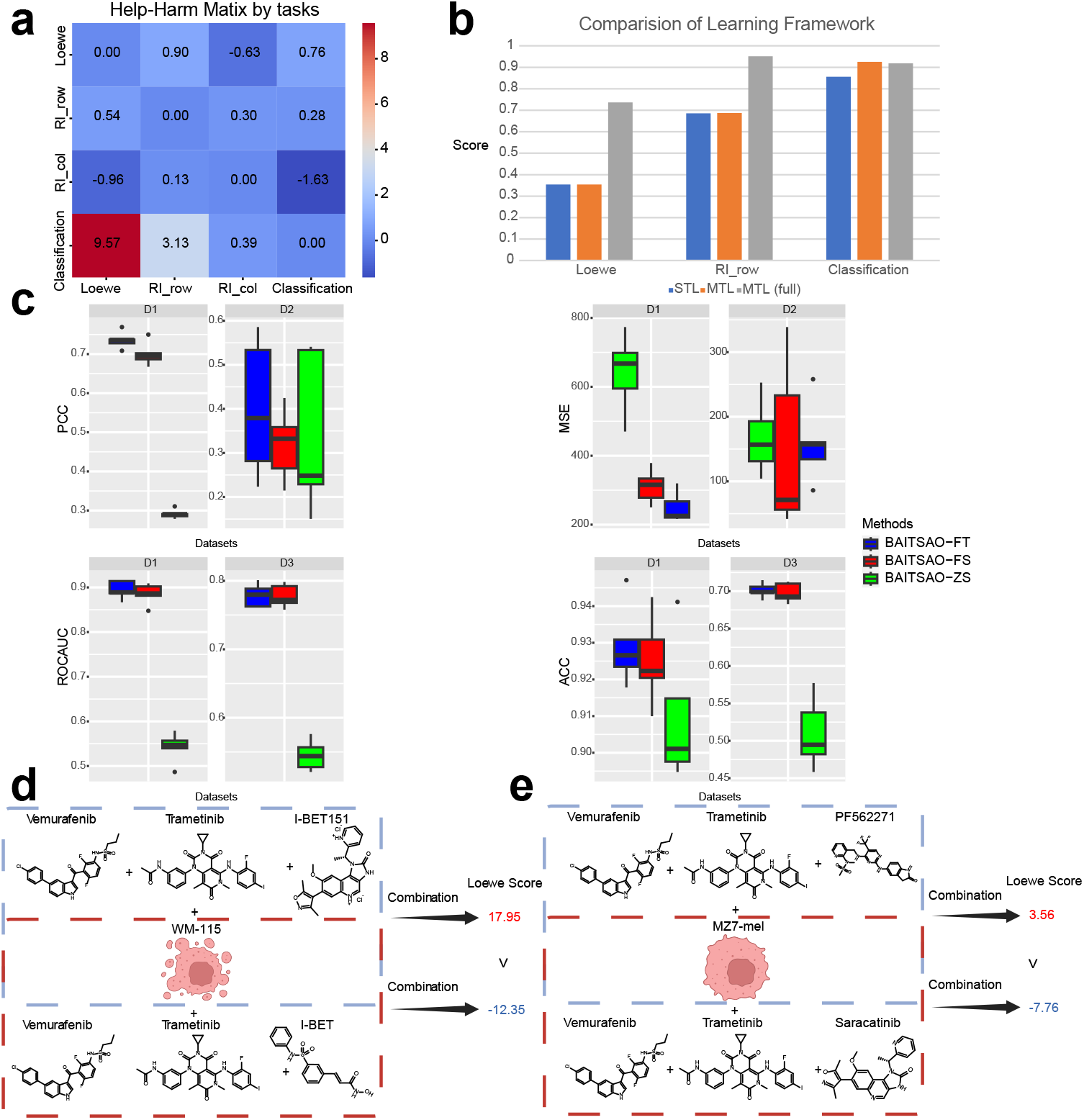
Results under the multi-task learning framework. (a) The Help-Harm matrix for different combinations of tasks. The values indicate the percentage (unit: %) of improvement using multi-task learning compared to single-task learning (STL) defined by the tasks in rows. The columns represent the paired tasks. We boldfaced blocks with increments larger than 0.5%, which is a threshold reported in [70] as acceptable improvement and half of the natural threshold 1%. (b) Comparisons for the results under MTL and STL. The metric for regression tasks, including Loewe and RI row, is PCC. The metric for the classification task, including Classification, is ROCAUC. (c) Comparisons for the results under different training settings. Data are presented in boxplots (n=5 per group; center line, median; box limits, upper and lower quartiles; whiskers, up to 1.5 *×* interquartile range; points, outliers). Here *BAITSAO-FT* represents that we fine-tuned the pre-trained model, *BAITSAO-ZS* represents that we applied the pre-trained model for these tasks under zero-shot learning framework, and *BAITSAO-FS* represents that we did not use the pre-trained weights for these tasks. Here FT means fine-tuning, ZS means zero-shot learning and FS means from scratch. We included four metrics across three datasets for comparisons. (d) The first example of tri-drug cases for drug synergy prediction with BAITSAO. (e) The second example of tri-drug cases for drug synergy prediction with BAITSAO. Source data are provided as a Source Data file.

We then predicted the synergistic effect for the combination of three drugs (tri-drugs) and one cell line, with two examples shown in Figures 5 (d) and (e). The drug names and cell line names were extracted from DrugCombDB [71], which did not provide the observed synergistic information for the existing combinations. To enhance the reliability of our prediction results, we relied on Monte Carlo Dropout (MC Dropout) [72, 73] and ran inference 100 times to generate the prediction interval of different drug combinations. According to [74], MC Dropout was the only method considered in this benchmarking paper to estimate the mean and variance without extra hyper-parameters. Our full prediction results are summarized in Supplementary file 3. Here we compared the difference between the two combinations by changing the third drug. We found that the combination with I-BET151 was predicted to have a positive sign in the synergy score under Loewe, while the combination with I-BET was predicted to have a negative synergistic effect. As an explanation, although these two drugs can both combine with bromodomain and extra terminal domain (BET) with the same major targeted proteins [75, 76], I-BET151 was reported as an optimized version with excellent BET target potency and selectivity [76]. Therefore, we expected I-BET151 to have better efficacy and thus a higher synergy score. Another example from Figure 5 (e) presents the difference between PF562271 [77] and Saracatinib [78] as a third drug under the cell line MZ7-mel. The combination with PF562271 had a higher predicted synergy score compared with Saracatinib, which was supported by the experimental results from [79] as PF562271 generated higher growth inhibition. Therefore, the results from BAITSAO can help researchers to optimize drugs with higher synergistic effect and better clinical outcomes.

### Sensitivity analysis

Here we investigated the sensitivity of model training based on the statistics we collected. Figure 6 (a) displays the ablation results by considering different types of embeddings as well as different types of combination rules for embeddings as model input. *BAITSAO* denotes our final choice for pre-training and fine-tuning. *BAITSAO-v3* denotes that we utilized the updated embeddings from OpenAI in 2024 [80]. *Mean* denotes that we took the mean of drug embeddings and cell embeddings as input for training. *Sum* denotes that we took the sum of drug embeddings and cell embeddings as input for training. *SentStack* denotes that we stacked the descriptions of different drugs and used the modified description to generate drug embeddings, and then stacked such drug embeddings with cell-line embeddings. *Stack* represents that we stacked the drug embeddings and cell embeddings by rows. *Rdkit* [41, 42] represents that we generated embeddings from Rdikt with SMILES and stacked the embeddings with cell embeddings from LLMs. This figure shows that averaging the drug embeddings and stacking them with cell embeddings by rows generated the best performance for all tasks. These results suggest the most effective way to incorporate embeddings from different sources to construct the datasets for training and testing. Moreover, our approach strikes a good balance between efficiency and performance. According to Figure 6 (b), the running time of BAITSAO without pre-training was significantly lower than the classical methods DeepSynergy and SVM for drug synergistic effect prediction. Moreover, the pre-trained BAITSAO with the fine-tuning framework converges at a much faster rate, thus pre-trained BAITSAO achieved an even faster running speed by comparing with MARSY and DeepDDs. Therefore, our training framework strikes a good balance between runtime and model performance. Both pre-training and fine-tuning stages can be finished with only one GPU, presenting no hardware barrier to deploy BAITSAO.

**Fig. 6.**
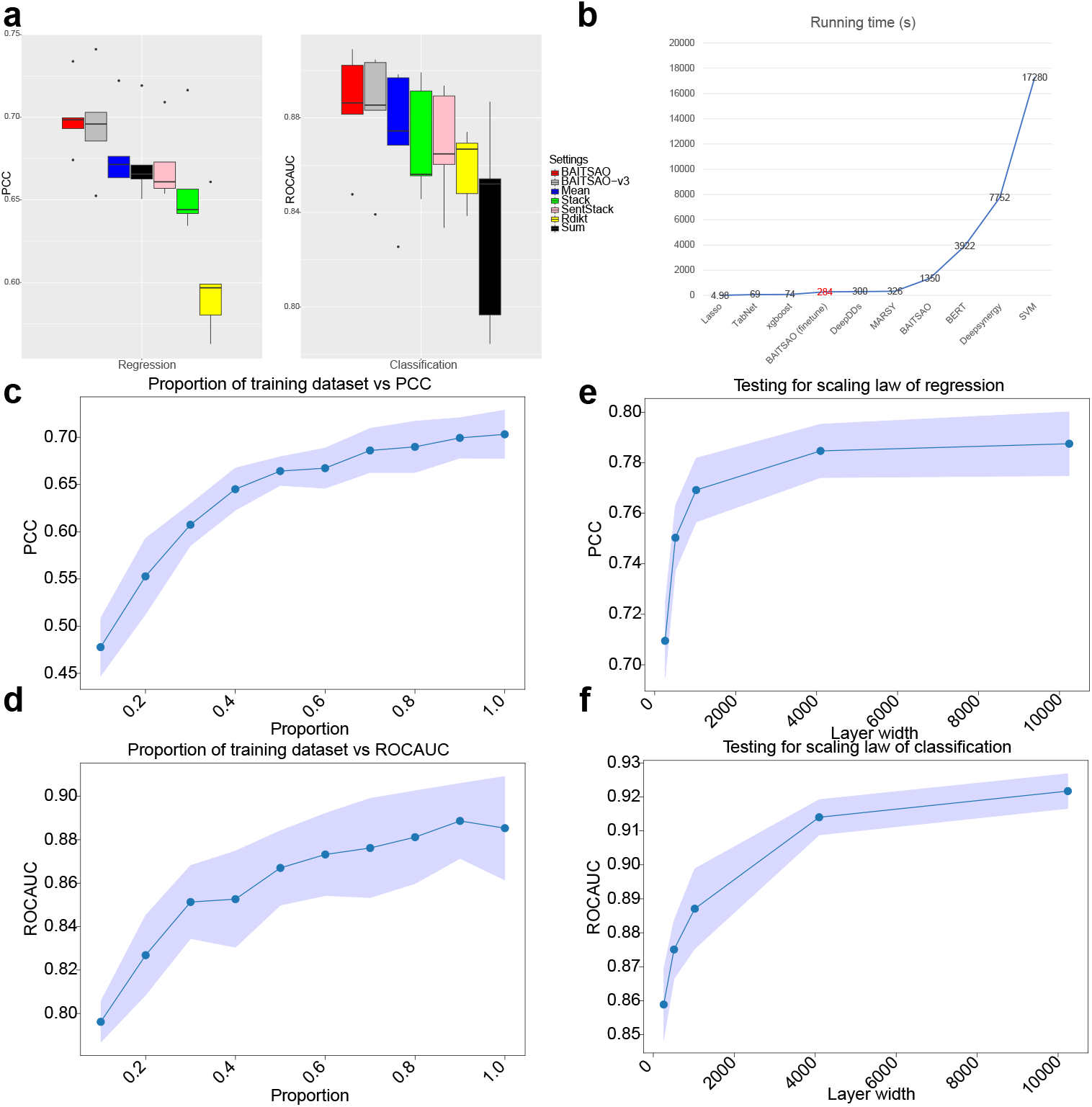
Statistics of model training. (a) Ablation test results for BAITSAO with different input formats. (n=5 per group; center line, median; box limits, upper and lower quartiles; whiskers, up to 1.5 *×* interquartile range; points, outliers). (b) The comparison of running time for different methods. We highlight the running time of BAITSAO and use the regression task as an example. (c) Plot for the proportion of training dataset and PCC for BAITSAO under the regression task. We reported (*µ ™ σ, µ* + *σ*) for each proportion, where *µ* represents the mean and *σ* represents the standard deviation. (d) Plot for the proportion of training dataset and ROCAUC for BAITSAO under the classification task. We reported (*µ ™ σ, µ* + *σ*) for each proportion. (e) Plot for layer width and PCC for BAITSAO under the regression task. We reported (*µ ™ σ, µ* + *σ*) for each proportion. (f) Plot for layer width and ROCAUC for BAITSAO under the classification task. We reported (*µ ™ σ, µ* + *σ*) for each proportion. Source data are provided as a Source Data file.

We performed ablation tests for the MTL strategy, shown in Supplementary Figure 12 for ablation of methods and Supplementary Figure 13 for ablation of task-specific layers. We compared the gradient matching-based approaches including PCGrad [81], GradVac [82], CAGrad [83], Nash-MTL [84] and the linear MTL framework LinearMTL [85, 86] with our revised UW approach and found that our choice generally had comparable or better results, especially for the classification task. Moreover, LinearMTL performed much worse than deep learning based methods on the regression type tasks. Therefore, we chose the revised UW as the method for the pre-training stage. Moreover, our final choice with one task-specific layer for each task had the best overall performance, and increasing the number of layers required more computing resources, thus we chose our design shown in Figure 1 (c).

We also analyzed the relation between the size of the training dataset and model performance. We adjust the proportion we used for model training and visualize the relation between proportion and metrics in Figures 6 (c) for regression and (d) for classification. From these figures, a larger proportion tended to increase the model performance, with its limit for proportion ≥ 0.9 for these two tasks. Moreover, using only 0.1% training dataset to train a model for a classification task can still generate relatively high ROCAUC, thus the classification task may not be difficult for BAITSAO.

In Figures 6 (e) and (f), we examined the scaling law [87, 88] of BAITSAO. We adjusted the layer width of our model and plotted the relation between the layer width in the hidden layer and model performance for the regression task and the classification task. These figures show that we can model the relation between model parameters and model performance to predict performance, where more parameters lead to better performance. Therefore, the performance improvement of our model with scaling can be explained by the scaling law, and our model has good scalability. Our findings can help us understand the model training process in a better approach and determine the optimized source allocation of a fixed compute budget. For example, for machines that cannot support the version of BAITSAO with layer width as 10240, the version of BAITSAO with layer width as 4096 can also have acceptable performances and can be considered to deploy.

## 3 Discussion

Predicting drug synergistic effect is important for drug development and patient treatment. In the past, limited by available experimental data, information on drugs/cell lines, and pipelines to predict drug synergistic effect, there are few approaches to predicting drug synergistic effect for general use. With the help of large-scale drug synergy information databases, LLMs, and an MTL framework, we introduced BAITSAO as a unified model with a general pipeline for drug synergistic effect prediction as well as single-drug inhibition prediction. BAITSAO optimized the network architecture through comprehensive benchmarking analysis and was pre-trained based on the latest large-scale databases. It achieved top-tier performance in both regression tasks and classification tasks for drug synergistic effect prediction.

There are two major contributions of our work. Firstly, we presented a unified pipeline to construct datasets for synergistic effect analysis for both drugs and cell lines based on the embeddings from LLMs, thus we mitigated the difference caused by aliases for drugs and cell lines of different datasets. We demonstrated that the embeddings contained functional information for drugs and cell lines. We proposed a new design to construct training datasets, thus we only need to utilize the overlapped information across datasets for drug synergy analysis. Secondly, we pre-trained a unified model with a MTL framework for drug synergy analysis and single-drug inhibition analysis supported by rigorous task-selection steps. We demonstrated that BAITSAO benefited from the pre-training process and had good generalization ability with fine-tuning in fewer steps compared with the training process from scratch. Moreover, pre-trained BAITSAO showed its potential as a good zero-shot reasoner for drug synergy prediction under the classification settings. Therefore, we overcame the generalization issue in previous work based on transfer learning [21] and proposed a new avenue for the construction of BAITSAO for drug synergy analysis.

We conducted a sensitivity analysis to offer guidance for future model deployment. We showed that our current hyper-parameter settings and data construction methods are the optimal choices by hyper-parameter tuning and ablation tests. We also analyzed the relation between the proportions of data we used for training and model performance. While increasing training data proportions tended to improve prediction, BAITSAO performed well for the classification task for small data scales. Finally, we investigated the scaling law of BAITSAO and showed that the model performance is predictable and we could increase the model performance by scaling up BAITSAO for drug synergy prediction.

In conclusion, we have developed BAITSAO, an explainable model for drug synergy prediction, and demonstrated the superiority of BAITSAO over other methods by comprehensive benchmarking analysis and rigorous sensitivity analysis. We hope that BAITSAO can help researchers to better understand the process of drug synergistic effect prediction and further help in optimizing drug structures for drug design and discovering novel drug combinations with synergistic effects for clinical usage.

Furthermore, we also found that BAITSAO might not work well for drugs without a clear functional or chemical description in the early stage of drug development, which is a potential limitation of our application scenarios of all functional-based synergy predictors. In the future, we plan to incorporate more updated drug synergy databases to keep this model updated, and we also plan to combine this model with information from genomics including single-cell data [89] and genome-wide association studies (GWAS) [90], especially for early-stage and novel drugs.

## 4 Methods

### Problem definition

In this manuscript, we intend to construct a dataset 𝒟 = (*X, Y*) and pre-train a model known as ℳ for the prediction of values in *Y* ^*n*×*t*^, where *n* represents the number of combinations between drug pairs and the cell line, and *t* represents the number of tasks. Here *X*^*n*×*p*^ represents the feature space with *n* samples and *p* features. We then split the dataset 𝒟 into 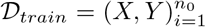 for training and 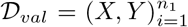 for validation. Our target is to train a model ℳ^∗^ based on 𝒟_*train*_ and then select the optimal model based on 𝒟_*val*_. That is,

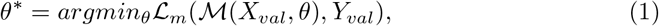

where ℳ (, *θ*) represents the pre-trained model with parameter *θ*, and *θ*^∗^ represents the optimal model parameters. ℒ_*m*_ represents multi-task learning loss. After obtaining the optimal model, we apply the model ℳ^∗^(, *θ*^∗^) for a new dataset containing out-of-distribution (OOD) data, known as 𝒟_*test*_.

### Construction of pre-training datasets and testing datasets

One major contribution of our work is to unify the features we need to predict the drug-related information for both the synergistic effect and inhibition effect. We at least need the names of drugs and cell lines. Considering we have a drug pair (*d*_1_, *d*_2_) and a cell line (*c*_1_), our idea is to generate the description of both drugs as *W* (*d*_1_), *W* (*d*_2_) and the cell line as *W* (*c*_1_) based on LLMs such as GPT 3.5, and then utilize the embeddings tool of GPT 3.5 to transfer the text description into embeddings with *e* dimensions. Therefore, our final sample *x* ∈ *X* is defined as:

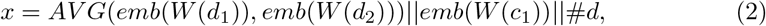

where *AV G*() represents the functions to compute the mean of the given variables, and *emb*() is the function to obtain the embeddings of the input. #*d* represents the number of drugs we used, which can be encoded as embeddings [91]. We take the unbiased estimation of the drug combination in the feature levels by computing the average value of embeddings, and we show that this approach works better than other types of feature integration in the sensitivity analysis section of the manuscript. Notably, this approach also scales for more drug combinations. Considering the case of *k* drugs with the cell line *c*_*i*_, we define one sample *x* ∈ *X* as:

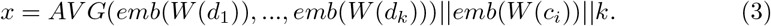

Therefore, for an arbitrary input dataset with feature space *X* containing drug information and cell-line information, we can transfer the samples in the given dataset from the text space to the numerical space, thus we unify the input data format for this task. Furthermore, to predict the drug synergistic effect, we consider both regression and classification. In the case of regression, we intend to predict the specific synergy score of samples. To compute the synergistic effect based on IC 50 information, under different rules, we can have different scores. Here we consider four methods to model the synergy scores, known as HSA, Bliss, Loewe and ZIP. If we consider *N* drugs with multi-drug combination effect as *E*_*A,B*,…,*N*_ and we intend to compute the synergy scores *S*_*HSA*_, *S*_*Bliss*_, *S*_*Loewe*_, and *S*_*ZIP*_, according to [5], we have:

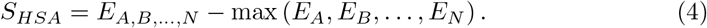

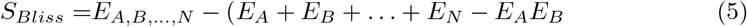

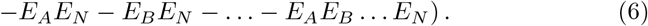

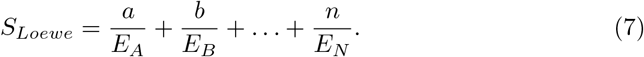

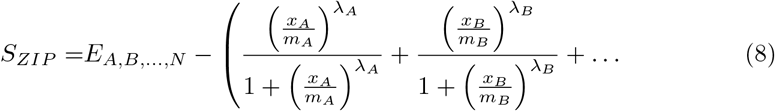

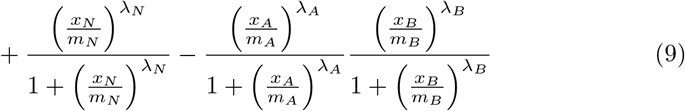

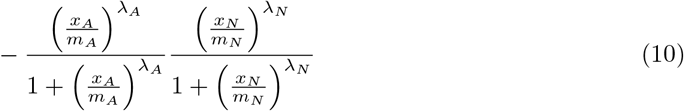

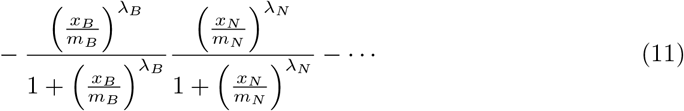

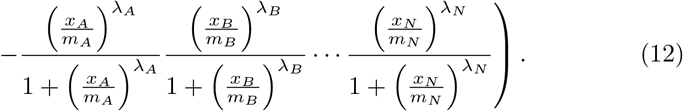

Here we have *E*_*A*_, *E*_*B*_, …, *E*_*N*_ as measured responses of different drugs, and *a, b*, …, *n* represent the doses of the single drugs we need to produce the combination effect. Moreover, to compute *S*_*ZIP*_, we have *x*_*N*_ as the dose of drug *N* fitted with the four-parameter log-logistic model, and *m*_*N*_ represents the dose we need to produce the half-maximum effect (IC 50). We also have *λ*_*N*_ as the shape parameter to indicate the slope of the dose-response curve. In the MTL case, we consider *S*_*Loewe*_ for the targets of regression. We also pre-train models to predict the other three scores. All of the synergy scores are extracted from the database of DrugComb.

In order to characterize the inhibitory effects of individual drugs, we introduce the RI score in the prediction task. RI score is the normalized area under the *log*_10_ − transformed dose-response curves. RI scores of all drugs are also extracted from the database of DrugComb.

In the case of classification, we intend to predict whether the given drug pair has a synergistic effect under a specific cell line, which is a binary classification problem. To construct the dataset for this task, we set the threshold of *S*_*Loewe*_ to binarize the synergistic effect of different drug combinations. Since not all of the testing datasets in the real world contain data for both regression and classification, thus introducing a classification task is meaningful.

### Investigation of embeddings

We set up different methods to ensure that embeddings from LLMs contain the necessary information to describe the properties of drugs and cell lines. We consider two prompt engineering approaches for description generation, including MetaPrompt [47] and Chain-of-Thought (COT) [47]. MetaPrompt introduces a system prompt for LLMs and generates the outputs conditioned on the context. COT allows LLMs to obtain complex reasoning capabilities by forcing models to address the problem with intermediate steps. We also generate text descriptions and embeddings for drugs and cell lines from the dataset used by Deep-synergy (D1). We check the correctness of the all descriptions and the similarity of 10 sampled embeddings across different drugs to evaluate the correctness. Moreover, we change the random seed to generate different descriptions as well as embeddings to check the variance of drug embeddings from the same drug. We also record the description of cell lines in Supplementary file 1. Furthermore, we modify CPA to predict gene expression under different perturbations enhanced by drug embeddings from LLMs. In this step, we replace the original drug embeddings used in the CPA with our new embeddings. This approach allows us to check the correctness of embeddings from the application perspective. We use *R*^2^ scores to evaluate the performances. To compute the *R*^2^ score, we have the ground truth synergy score *y* and predicted synergy score *ŷ* and follow its definition:

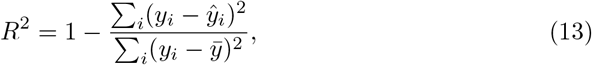

where 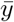 means that we compute the average value of the input variable *y. R*^2^ represents the explanation of independent variables for the dependent variable. Higher *R*^2^ means better model performance.

Therefore, our assessment of the quality of embeddings takes into account meanings, variance, and applications.

### Hyper-parameter searching

We summarize the search space for hyper-parameters of each method in Table 1. The best hyper-parameter setting is determined by the performance of models based on the validation dataset.

**Table 1.** Hyper-parameter search space for each method ranked in alphabetical order.

| Methods | Searching space |
| --- | --- |
| BAITSAO | lr:[1e-5,1e-4]; Dropout:[0.1,0.3] |
| BERT | epochs:[5,10]; lr: [1e-5,1e-4] |
| DeepDDs | Optimal hyper-parameters from [19] |
| DeepSynergy | Optimal hyper-parameters from [14] |
| Lasso | alpha:[1,10] |
| MARSY | Optimal hyper-parameters from [18] |
| SVM | C:[1,5] |
| TabNet | n_steps:[1,5]; n_a:[8,64]; n_d:[8,64]; gamma:[1.1,1.5] |
| TreeComb | max_depth:[10, 100]; n_estimators:[10,50]; min_child_weight:[1, 3] |

Here lr means learning rate, Dropout means dropout rate (the ratio of neurons we intend to close during the training process), max depth means the maximal depth for tree-based models, n estimators means the number of estimators for tree-based models, min child weight means the minimal weights of child nodes in tree-based models, C means the regularization weight for SVM, n steps means the number of decision steps in the model architecture, n a means the width of the attention embedding for each masked choice, n d means the width of the prediction layer, gamma means the coefficient for the feature re-usage in the masking process, epochs mean the number of epochs we used to train the model, alpha represents the regularization coefficient for Lasso. We present the results under different hyper-parameters for BAITSAO in Supplementary Figures 14 (a) and (b). We find that lr plays a more important role in the training process while adjusting the dropout rate does not affect the model performance much.

### Selection of model architecture

After setting up the pre-training dataset, we seek a suitable model architecture. Since deep neural networks (DNNs) related methods have shown impressive performance as a base model for large-scale models [92, 93], we construct the pre-training architecture of BAITSAO based on DeepSynergy. To assess the strength of our model architecture, we remove the pre-training step and compare BAITSAO with other methods for both the regression task and classification task with three different datasets. We also determine the hyper-parameters of model training in this stage. The superiority of BAITSAO is shown in the Results section, and we expect to see its similar performance at both the pre-training and fine-tuning stages.

In the model architecture selection stage, we utilize Adam [94] as the optimizer and ReduceLROnPlateau [91] as the learning rate scheduler. The starting learning rate for D1 and D3 is 1e-5, while it is 1e-4 for D2. The dropout rate is 0.2, and the patience for the scheduler is 10. Our patience for early-stopping step is 100, and the maximum number of epochs is 1000.

### Explainability

The design of BAITSAO allows us to characterize the relevance between the specific gene and drug combinations across different cell lines. To perform this analysis, the input format of one combination becomes:

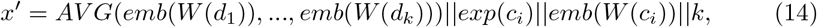

where *exp*(*c*_*i*_) represents the gene expression profile for the cell line *i*. After the training process, we can extract the importance of different genes based on SHAP. For gene *j*, its importance for the synergistic effect of drug combinations (*d*_1_, …, *d*_*k*_) for cell line *c*_*i*_ can be calculated as:

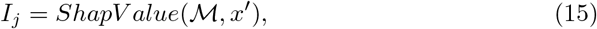

where *I*_*j*_ represents the importance and *x*^′^ was defined above. *ShapValue*() is a function to compute the importance with model ℳ and input *x*^′^. Here larger *I*_*j*_ represents more importance in the prediction process.

We select 1000 highly-variable genes for the analysis of explainability. This number is determined by adjusting the number of genes to achieve the best model performance. Our tuning results are shown in Supplementary Figure 15.

For the bulk RNA-seq datasets of different cell lines, we use DESeq2 to identify DEGs by comparing groups with and without predicted drug synergistic effects.

For the two scRNA-seq datasets used for validating our selected genes, we follow the pre-processing pipeline of Scanpy [95] and run the Wilcoxon rank-sum test to access the list of DEGs.

### Pre-training BAITSAO under the multi-task learning framework

Here we explain our settings for the multi-task learning framework. After filtering tasks based on the constructed help-harm matrix, we consider three tasks: 1. Prediction of *S*_*Loewe*_ as a regression task. 2. Prediction of RI for one drug as a regression task. 3.Prediction of drug synergistic effect as a binary classification task. Therefore, we have two regression tasks and one classification task, and their loss functions are represented as ℒ_1_, *ℒ*_2_, and ℒ_3_. Traditionally, we construct the final loss function as a linear combination:

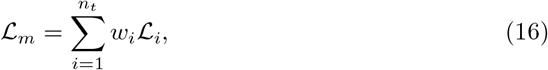

where *w*_*i*_ represents the pre-defined weights for the loss function ℒ_*i*_ and *n*_*t*_ = 3. However, determining the values of the weights is difficult. Moreover, it is a strong assumption that the weights do not change during training is also a very strong assumption. Therefore, we introduce the uncertainty of loss function in this process and make the weights learnable. Typically, by choosing mean squared error (MSE) as the loss function in the training process for the regression task, we have the equivalent maximum likelihood framework of a Gaussian distribution for prediction output *y* and model ℳ. Therefore, the log-likelihood of the regression task can be represented as:

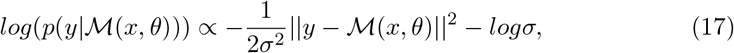

where the uncertainty is defined as *σ*, and *x* represents model input. *σ* is a learnable parameter. Similarly, for a classification problem, we can represent the log-likelihood based on a Softmax function, that is:

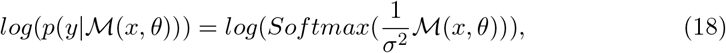

where *Softmax*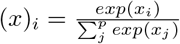, and *p* represents the length of *x*.

Therefore, the final loss function can be represented as maximizing the joint distribution of three tasks. In the validation stage, we minimize the maximal term in the loss function group rather than the original weighted loss function design from UW. We also add a constant term *ϵ* to ensure the numerical stability, so our final loss function is:

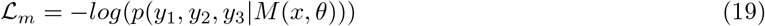

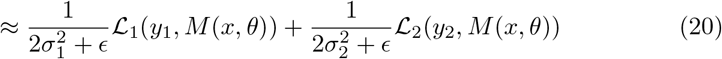

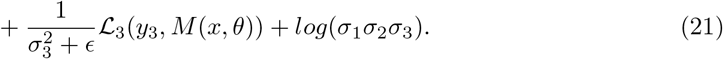

In the pre-training stage, we utilize Adam [94] as optimizer and ReduceLROnPlateau [91] as the learning rate scheduler. The starting learning rate is 1e-4, the dropout rate is 0.2, and the patience for the scheduler is 100. Our patience of early-stopping step is 500, and the maximum number of epochs is 1000. The number of combinations we used for pre-training is 739,652, including 4268 unique drugs and 288 unique cell lines.

After finishing the pre-training step, we test the model performance on the testing datasets under both the zero-shot learning case and the fine-tuning case with a parameter-freezing design. We also extend the prediction of the synergistic effect to the case of *n* (*n* ≥ 3) drug combinations. Finally, we include a tutorial in our code repository for both the fine-tuning approach and the zero-shot inference approach.

We also pre-train other baselines, including DeepSynergy [14], DeepDDs [19], and MARSY [18], based on the same dataset. Details of model comparison are discussed in the Results section.

### Zero-shot query and multi-drug prediction

Our model is capable of zero-shot synergy effect prediction. By transferring the knowledge and information of drugs and cell lines into embeddings through GPT 3.5 and the embedding layer, users can generate embeddings of arbitrary combinations as input for querying the synergy effects with a pre-trained BAITSAO.

For the combinations with three or more drugs, we directly generate the synergy score under the pre-trained model with the zero-shot learning framework. The three-drug case we used in the main text is from a known database, while it is possible to explore combinations with a larger number of drugs as long as the combinations are practical and meaningful. To access the determined predicted value, we do not use the dropout layers in the testing process. To access the predicted value with uncertainty, we keep the dropout layers in the testing process and repeat the prediction process for 100 times to access the estimation of mean and standard deviation for each combination. Such approach is known as MC Dropout.

We summarize the details of zero-shot query as a tutorial in our code repository.

### Model evaluation

We consider four different metrics to evaluate the performance of different models for the drug synergistic effect prediction task, with two metrics for regression and two metrics for classification.

For the regression task, we consider two metrics: Pearson correlation coefficient (PCC) and Mean Squared Error (MSE).

1. PCC: Since we know the ground truth synergy score *y* and predicted synergy score *ŷ*, we can directly compute the PCC as:

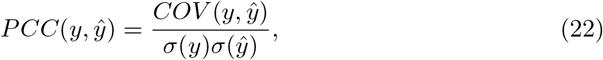

where *COV* () is the function to compute the covariance of two variables, and *σ*() is the function to compute the standard deviation of the input variable. Higher PCC means better model performance.
2. MSE: To compute the mean squared error, we have the ground truth synergy score *y* and predicted synergy score *ŷ* and follow its definition:

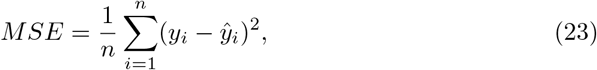

where *i* represents the index of samples, and lower MSE means better model performance.

For the classification task, we consider two metrics: Area under the ROC Curve (ROCAUC) and Accuracy (ACC).

1. ROCAUC: To compute this metric, we construct the relation between the truepositive rate and the false-positive rate under different probability thresholds. Such relation can be reflected in the ROC curve. We then compute the area under the ROC curve, and this area represents ROCAUC. Higher ROCAUC means better model performance.
2. ACC: To compute this metric, we have the ground truth synergistic effect condition *y*^*n*×1^ and predicted binary value *ŷ*, we then compute the ACC as:

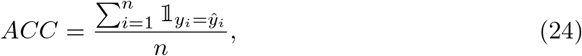

where 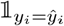 is a indicator function and only takes 1 when *y*_*i*_ = *ý*_*i*_. Higher ACC means better model performance.

We report the mean and standard deviation of these metrics by using five-fold cross-validation for each dataset.

### Overview of other methods

In this section, we summarize the benchmarking methods used in our work. These methods (ranked in alphabetical order) include:

- BERT: BERT is a pre-trained bi-directional transformer for language understanding. For this model, we construct the training datasets and testing datasets directly from drug descriptions and cell-line descriptions. The problem is then formalized as Question-answering case for both classification and regression tasks.
- DeepDDs [19]: DeepDDs is a Graph Neural Network (GNN)-based method for drug synergistic effect prediction. This method can only handle the classification task. The training dataset of DeepDDs is constructed base on features of drugs as graphs from chemical information and gene expression levels from cell lines.
- DeepSynergy [14]: DeepSynergy is a DNN-based method for drug synergistic effect prediction. This method can handle both the regression task and the classification task, by changing the loss function and the activation function of the last network layer. The training dataset of DeepSynergy follows its default mode, including features of drugs from chemical information and cell-line features from gene expression levels.
- Lasso [39, 52]: Lasso is a regularized regression method for drug synergistic effect prediction. This method can handle both the regression task and the classification task, by using the default mode or logistic regression mode with L1 penalty. The training dataset of Lasso is constructed based on the drug embeddings and cell-line embeddings from LLMs.
- MARSY [18]: MARSY is a DNN-based method with multi-task learning framework for drug synergistic effect prediction. This method can only handle the regression task. The training dataset of MARSY is constructed based on features of drugs from chemical information, gene expression levels from cell lines and tissue information.
- SVM [39, 49]: SVM is a machine learning method based on constructing decision-making boundaries for drug synergistic effect prediction. This method can handle both the regression task and the classification task, by using SVR or SVC. The training dataset of SVM is constructed based on the drug embeddings and cell-line embeddings from LLMs.
- TabNet [50]: TabNet is a DNN-based method with transformer architecture for drug synergistic effect prediction. TabNet combines the ideas from both neural network design and tree-model design. This method can handle both the regression task and the classification task, by changing the loss function and the activation function of the last network layer. The training dataset of TabNet is constructed based on the drug embeddings and cell-line embeddings from LLMs.
- TreeComb [15]: TreeComb is an explainable machine learning method based on XGBoost for drug synergistic effect prediction. This method can handle both the regression task and the classification task, by using XGBREGRESSOR or XGBCLASSIFIER. The training dataset of TreeComb is constructed based on the drug embeddings and cell-line embeddings from LLMs.

### Datasets preparation

We utilize public datasets from DrugComb v1.5 for pre-training. For the regression task, we have one dataset from DeepSynergy (as D1, which is processed in the original paper) using Loewe as the synergy score computation method. We also have one dataset from MARSY (as D2, which is processed in the original paper) using ZIP as the synergy score computation method. For the classification task, we have one dataset from DeepSynergy (as D1) using the Loewe synergy score with a threshold. We also have one dataset from DeepDDs with a known binary synergistic effect condition (as D3 [96]). For multi-drug synergistic effect inference, we utilize one dataset from DrugCombDB. Every dataset at least contains the names of drugs and cell lines.

## 5 Data availability

We do not generate new data in this research and all data used in this manuscript are publicly available. The DrugComb data used in this study are available under accession code [DrugCombDownload](https://drugcomb.org/download/). The training and testing data used in this study are available under accession code [D1](https://www.bioinf.jku.at/software/DeepSynergy/), [D2](https://github.com/Emad-COMBINE-lab/MARSY), and [D3](https://github.com/Sinwang404/DeepDDs/tree/master). The scRNA-seq data used in this study are available under accession code GSE215121 and SCP109. We collect the information of downloading training datasets as well as their statistics in Supplementary file 4. Source data are provided with this paper.

## 6 Reproductivity and codes availability

We used the resources from the Yale High Performance Center (Yale HPC) and UCLA Computing Servers to conduct all of the experiments. Our maximum running time for each dataset was 24 hours and maximum RAM was 100 GB. The version of GPU we used is NVIDIA A5000 (24 GB) for fine-tuning and single task learning, and NVIDIA A100 (40GB) for pre-training. The codes of BAITSAO can be found in https://github.com/HelloWorldLTY/BAITSAO and https://doi.org/10.5281/zenodo.15105815 with MIT license. The pre-trained weights can be found in https://huggingface.co/iLOVE2D/BAITSAO. The version of softwares used for data collection and model training is summarized in Supplementary file 4.

## 7. Acknowledgements

We thank Yijia Xiao for the fruitful discussion about training the model. We thank Wangjie Zheng, and Tianqi Chen for helpful discussion about designing drug embeddings. We thank Dr. Ning Sun, Xinyi Chen, Dr. Gefei Wang, Dr. Yingxin Lin and Chenyu Wang for the suggestions about approaches to improve the experimental design. We also thank Jiawei He for helping us prepare the molecular file. This project was partly supported by NIH grants U24HG012108 and P50 CA196530 awarded to Dr. Hongyu Zhao, and OpenAI Researcher Access Program and Google Cloud Research Program awarded to Tianyu Liu.

## 8 Author Contributions Statement

T.L. proposed this study. T.L., T.C., and X.L. designed the model. T.L. ran all the experiments. T.L., X.L. and H.Z. wrote the manuscript. H.Z. supervised this study.

## 9 Ethics declarations

### 9.2 Ethics and Inclusion

Although BAITSAO is not biased on gender, races, and other factors, the users are solely responsible for the content they generate with models in BAITSAO, and there are no mechanisms in place for addressing harmful, unfaithful, biased, and toxic content disclosure. Any modifications of the models should be released under different version numbers to keep track of the original models related to this manuscript. The users must comply with the laws of the country in which they are located.

The target of current BAITSAO only serves for academic research. The users cannot use it for other purposes. Finally, we are not responsible for any effects produced by other users.

## A Analysis of important genes based on synergy prediction and single-cell datasets for melanoma cells

In Figure 17, we show the explainability of different genes based on the drug combination: DEXAMETHASONE-DINACICLIB on the cell line A2058, which is a cell line for melanoma cells [97]. We repeated the training process three times with different random seeds and computed the average importance of genes for the analysis of a single cell line, thus we reduced the negative effect of randomness. We also found support for these two drugs as CDK inhibitors for affecting the expression levels of the 1st gene S100B [98, 99] and 2nd gene TYR [100, 101] ranked by the importance. Therefore, by analyzing the case of a single cell line, we linked the targets of individual drugs to genes, thus offering more information on treatments for the specific cancer type. Furthermore, we list the rank of selected genes based on their variance in Figure 17, and the top 5 genes have lower rank compared with important genes from all cell lines. Moreover, we considered running enrichment analysis based on GO and MsigDB for this set of genes, and the results are shown in Supplementary Figures 6 and (d). From these two figures, we found that the collected genes were significantly enriched in pathways related to melanosome, which also has high relevance with melanoma [102]. Moreover, compared to the analysis of multiple cell lines, we obtained fewer cancer-specific signals for the analysis based on the single cell line.

We also considered two approaches to validate the selected genes from cell line A2058 with the help of scRNA-seq datasets. We utilized one scRNA-seq dataset combined by [103, 104], known as the disease-control scRNA-seq dataset, to verify whether the selected genes are differentially expressed in malignant cells from patients. We analyzed its differentially expressed genes (DEGs) by running the Wilcoxon rank-sum test between the diseased cells and the control cells. We found that 16 out of the 20 genes were significant in the malignant cells. For four genes that were not significant, their ranks of expression levels across all genes are 13000+ by descending, so their contributions might be hidden by their measured expression levels in this dataset. Therefore, BAITSAO successfully captured the difference between tumor microenvironments and healthy cells.

Moreover, we utilized the other scRNA-seq dataset [105] labeled as acral melanoma (AM) and cutaneous melanoma (CM), known as AM-CM scRNA-seq dataset, to analyze its differentially expressed genes (DEGs) by running the test of the different diseased cases from patients. We show the UMAP plots of this scRNA-seq dataset in Supplementary Figures 18 (a)-(b). We found that 19 out of the 20 genes had low adjusted p-values, which meant that most of them were DEGs by comparing AM cells with CM cells, thus supporting the potential contribution of our model to uncover tumor heterogeneity.

We summarized our statistical analysis for these two datasets in Supplementary file 2. Our analyses therefore suggest that BAITSAO can provide information about potential targets of combined drugs for melanoma or other cancer types.

## B What are the advantages of BAITSAO comparing with other LLM-based methods?

In the area of drug synergy predictions based on Large language Models (LLMs), there are two existing methods, SynerGPT [25] and CancerGPT [26]. We discussed their limitations and differences from BAITSAO in the Introduction section of our submitted manuscript. Here we provide more details on the advantages of our method over these two other approaches by first summarizing the functions and challenges of these two methods in the Extended Data Table 1.

**Supplementary Tab. 1.** Comparison of SynerGPT and CancerGPT.

| Methods | Functions | Drawbacks |
| --- | --- | --- |
| SynerGPT | SynerGPT is a tool based on in-context learning. It models different examples containing drug pairs and one cell line based on their overlap to generate a synergy graph then uses the synergy graph as input to infer the synergetic effect of new examples. | 1. SynerGPT can only handle simple binary classification. 2. It does not consider prior biological knowledge. 3. It cannot handle multi-drug cases (with # of drugs≥3). 4. Its performance is close to DeepDDS according to their benchmarking results in Table 2. 5. It cannot be used to analyze gene-drug interactions and gene-gene interactions. 6. It is closed-source. |
| CancerGPT | CancerGPT is a tool based on pre-training and fine-tuning. It first pre-trains a model based on GPT-2 with biomedical context, and then incorporates the drug pairs and one cell line into the input sentence to form a question. The model's output, as the answer, will be treated as the prediction result. | 1. CancerGPT can only handle simple binary classifications. 2. It cannot handle multi-drug cases (with # of drugs≥3). 3. It cannot be used to analyze gene-drug interactions and gene-gene interactions. 4. It is closed-source. |

According to this table, both SynerGPT and CancerGPT have limitations that are addressed by BAITSAO, which is capable of using prior biological knowledge to predict synergetic effects for multi-drug combinations under both the classification and regression frameworks. According to our benchmarking results, BAITSAO outperforms baselines including DeepDDS, TreeComb, and MARSY significantly. Moreover, because SynerGPT and CancerGPT are not open-source (for CancerGPT we even cannot access their datasets for evaluation), we include BERT as a baseline to evaluate the performance of applying LLMs directly to address this problem, and BERT performs worse than BAITSAO. Moreover, we downloaded the datasets used by SynerGPT and performed prediction using BAITSAO, achieving an AUCROC score of 0.94, which is significantly higher than their reported score of 0.78. The results of our benchmarking analysis are shown in Figure 3. We are also willing to compare BAITSAO with SynerGPT and CancerGPT if they release their models in the future. Our method also allows researchers to explore gene-gene interactions as well as gene-drug interactions, which are not discussed in these two methods. Our method also introduces the idea of using embeddings from LLMs to analyze the similarity of drug functions. Finally, BAITSAO is not on a large scale (50M trainable parameters during the fine-tuning step), and it is fully open-source, which is friendly for the development of science and the extension from possible follow-up work.

### C Supplementary figures

**Supplementary Fig. 1.**
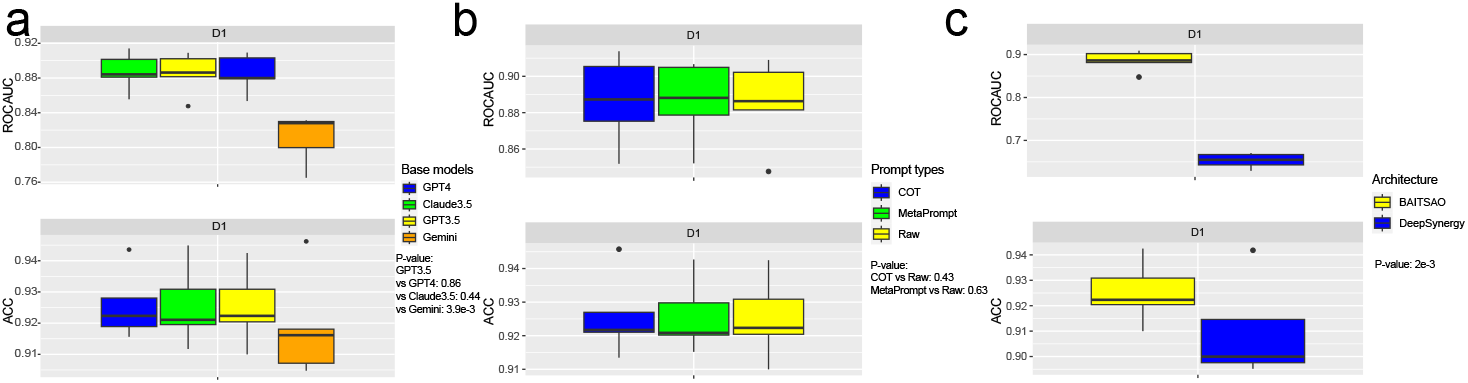
Evaluating the contributions of inputs and model architecture for drug synergetic effects. Data are presented in boxplots (n=5 per group; center line, median; box limits, upper and lower quartiles; whiskers, up to 1.5 *×* interquartile range; points, outliers). All of the experiments are performed based on dataset D1 and synergetic effect prediction is a classification task. The test is performed jointly for two metrics, based on Wilcoxon rank-sum test. (a) The performance comparison based on LLM embeddings from different base LLMs. (b) The performance comparison based on LLM embeddings from the drug descriptions generated by different prompts. (c) The performance comparison based on different model architectures.

**Supplementary Fig. 2.**
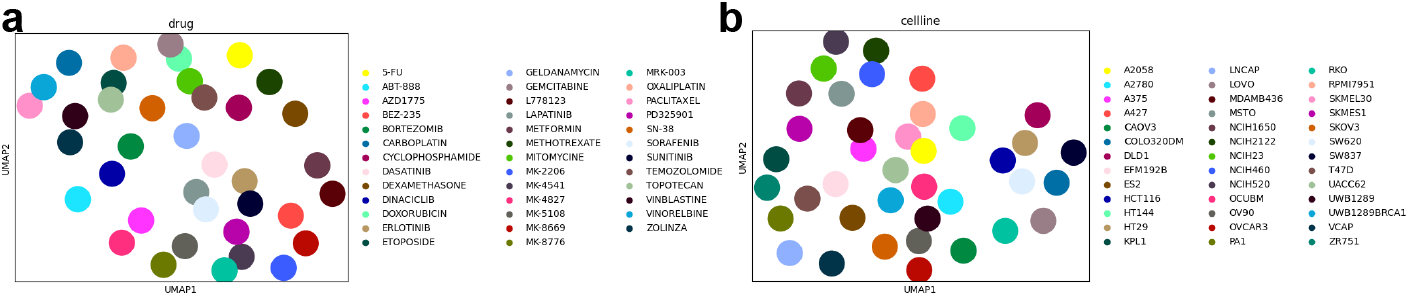
Visualization for the similarity of embeddings. (a) The UMAP plot for the drug embeddings from D1 colored by drug names. (b) The UMAP plot for the cell line embeddings from D1 colored by cell line names.

**Supplementary Fig. 3.**
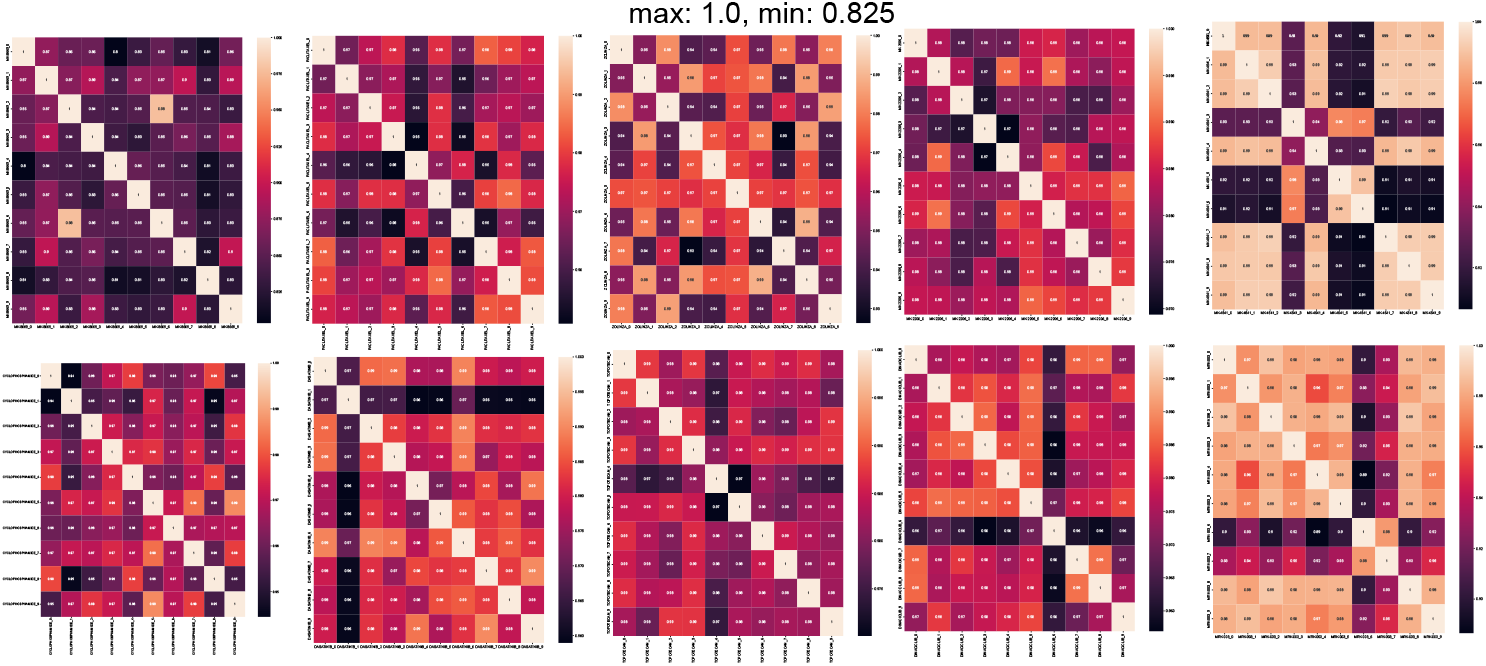
The group of heatmap for the similarity of embeddings under 10 random seeds of the 10 sampled drugs.

**Supplementary Fig. 4.**
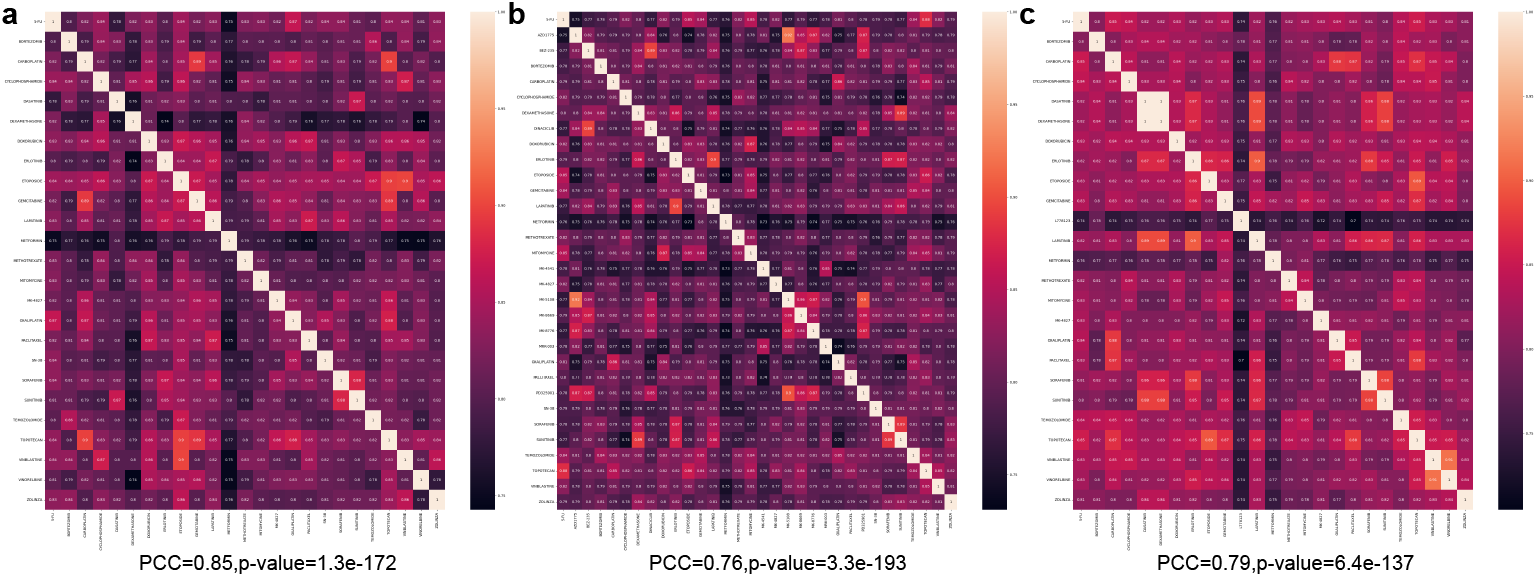
The group of heatmap for the similarity of embeddings from DrugBank under different drug properties. (a) The heatmap based on indication information. (b) The heatmap based on summary information. (c) The heatmap based on background information.

**Supplementary Fig. 5.**
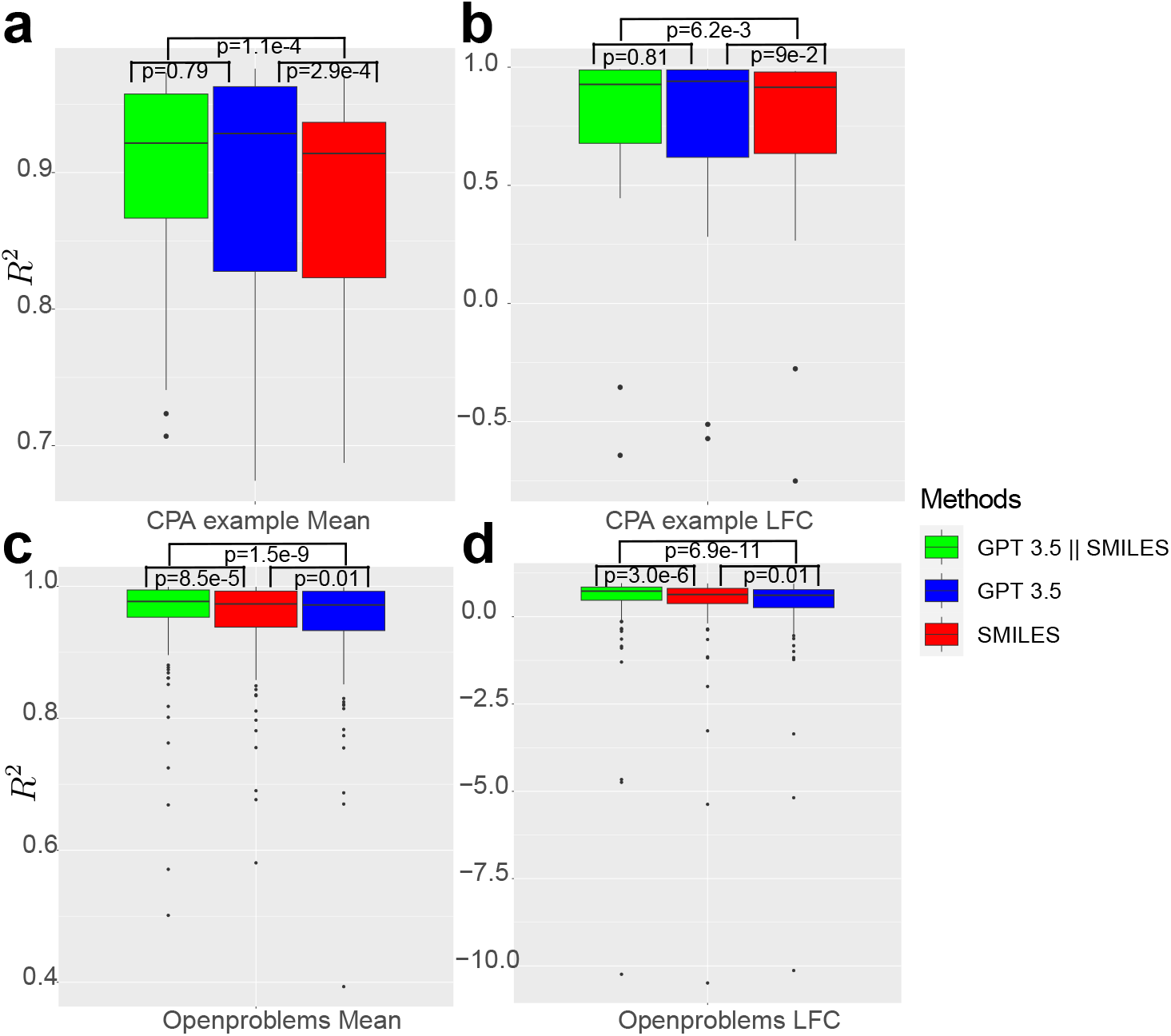
Exploration of drug-embeddings-augmented CPA results. Data are presented in boxplots (center line, median; box limits, upper and lower quartiles; whiskers, up to 1.5 *×* interquartile range; points, outliers). Higher *R*^2^ score means better performance. In panels (a)-(d), we also present the two-side p-values (p) based on Wilcoxon rank-sum test for each group. (a) *R*^2^ of the expression levels of deferentially expressed genes (DEGs) for the CPA example dataset colored by different settings (n=31). (b) *R*^2^ of the log-fold change (LFC) of DEGs for CPA example dataset colored by different settings (n=31). (c) *R*^2^ of the expression levels of DEGs for Openproblems dataset colored by different settings (n=137). (d) *R*^2^ of the LFC of DEGs for Openproblems dataset colored by different settings (n=137).

**Supplementary Fig. 6.**
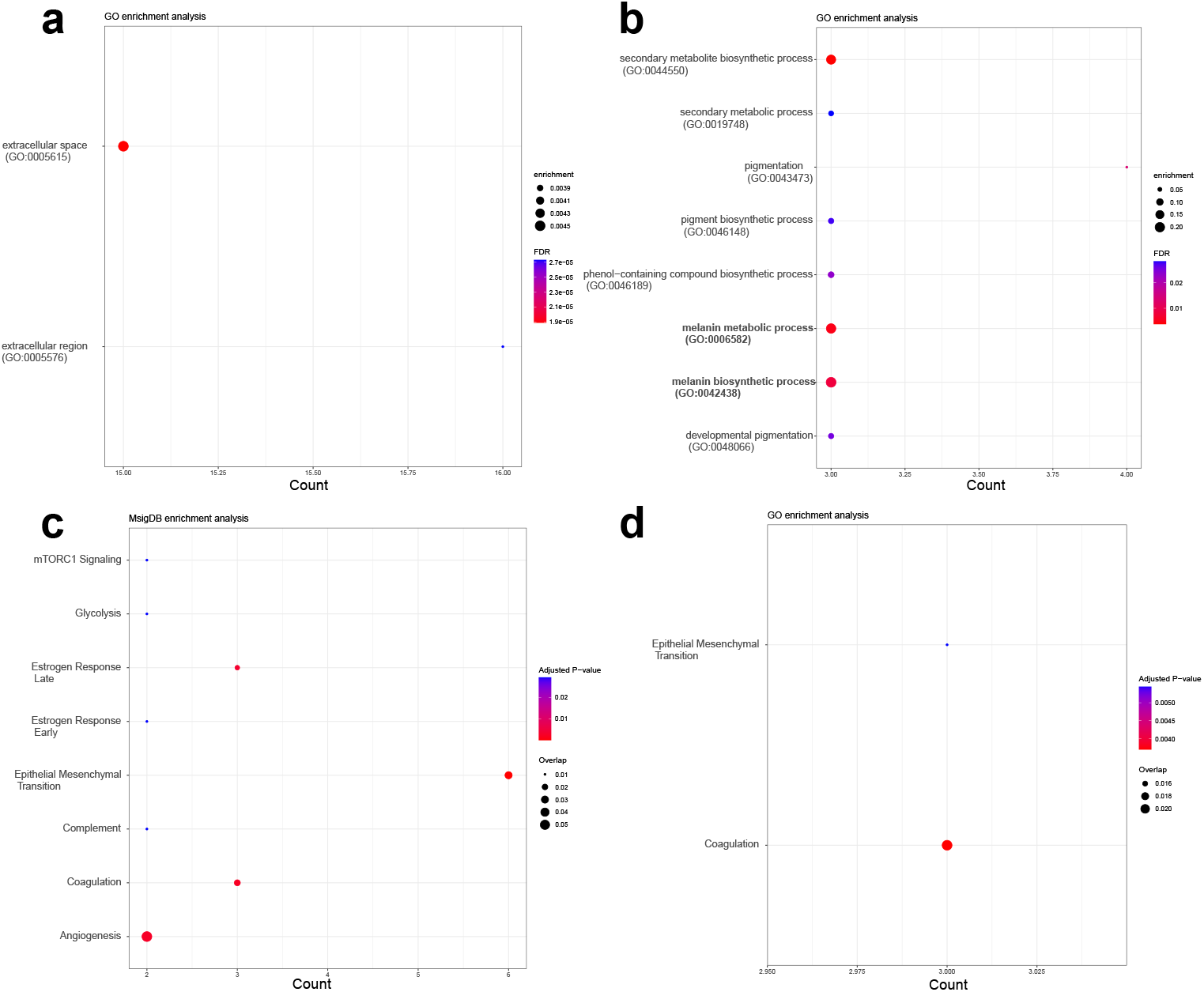
Results of gene enrichment analysis based on different databases. (a) Results of GO enrichment analysis using important genes from all cell lines. (b) Results of MsigDB enrichment analysis using important genes from all cell lines. (c) Results of GO enrichment analysis using important genes from the cell line A2058. We blodfaced the pathway related to melanosome. (d) Results of MsigDB enrichment analysis using important genes from the cell line A2058.

**Supplementary Fig. 7.**
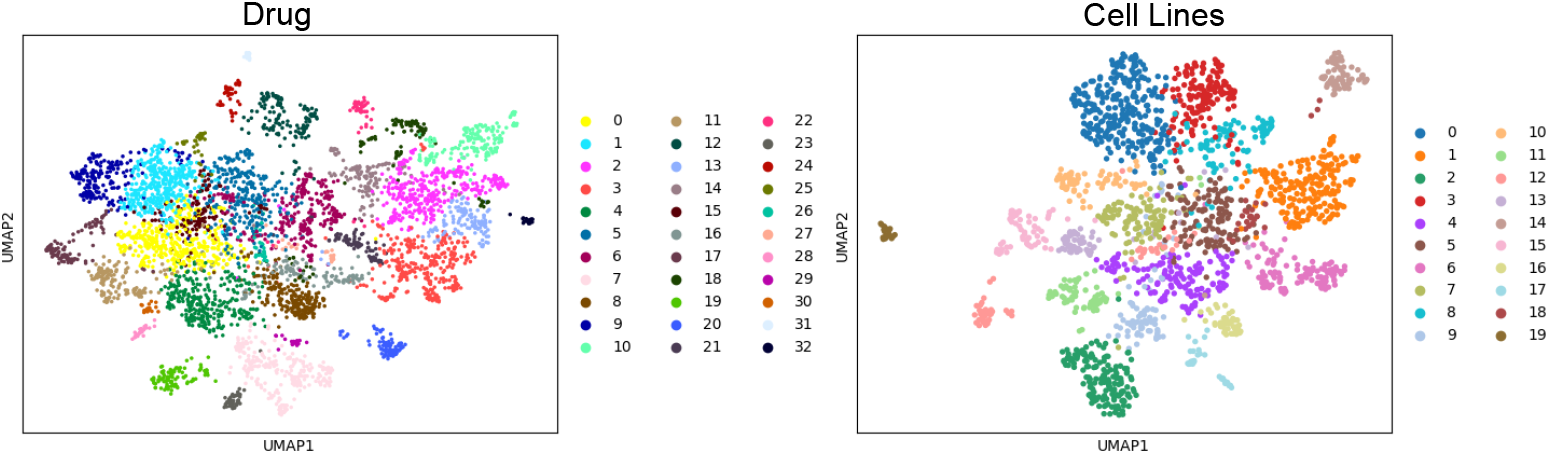
Visualization for the pre-training datasets. (a) The UMAP plot for the drug embeddings we used in the pre-training step, colored by Leiden clusters. (b) The UMAP plot for the cell-line embeddings we used in the pre-training step, colored by Leiden clusters.

**Supplementary Fig. 8.**
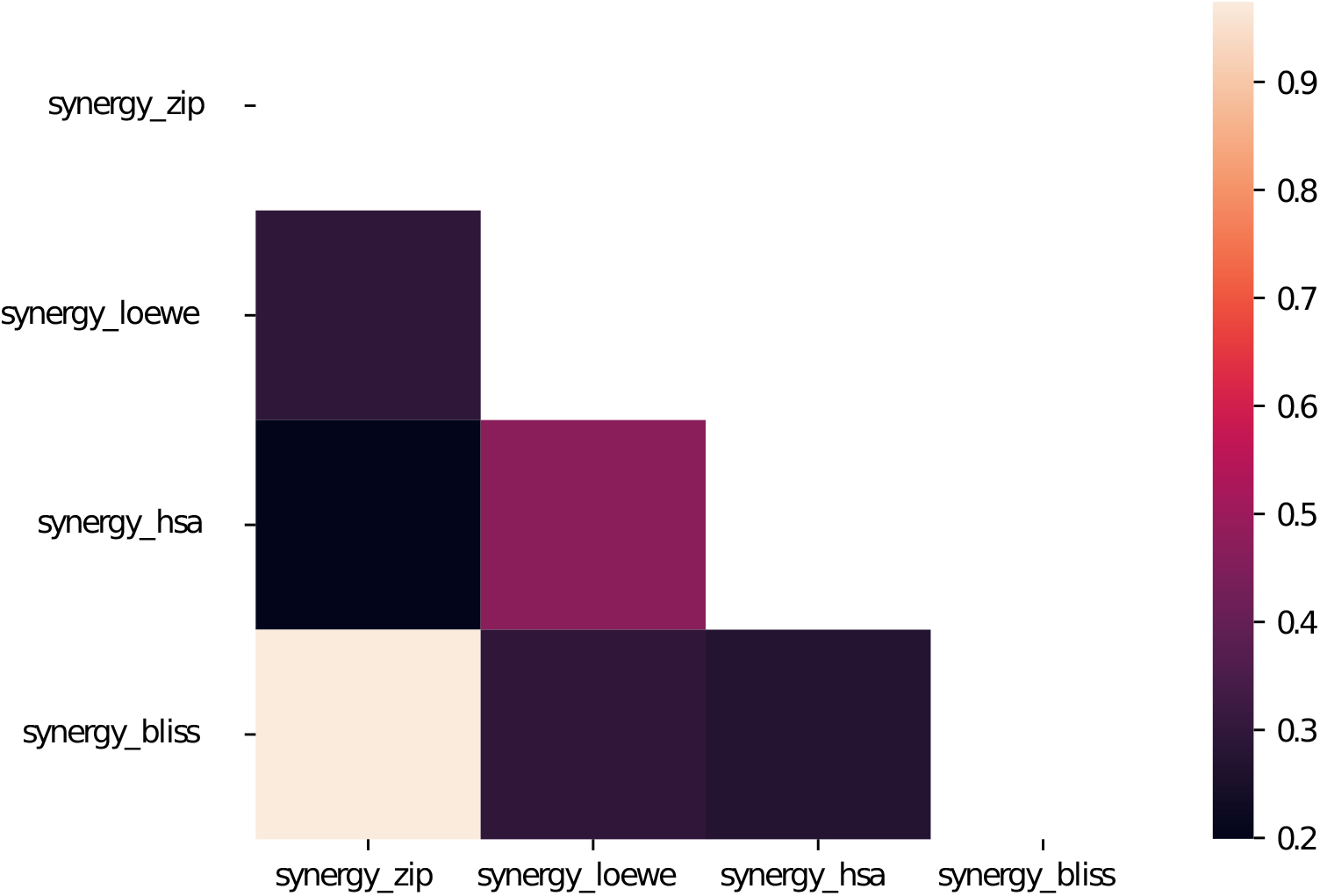
The heatmap for the PCC of different synergy scores from different computation methods.

**Supplementary Fig. 9.**
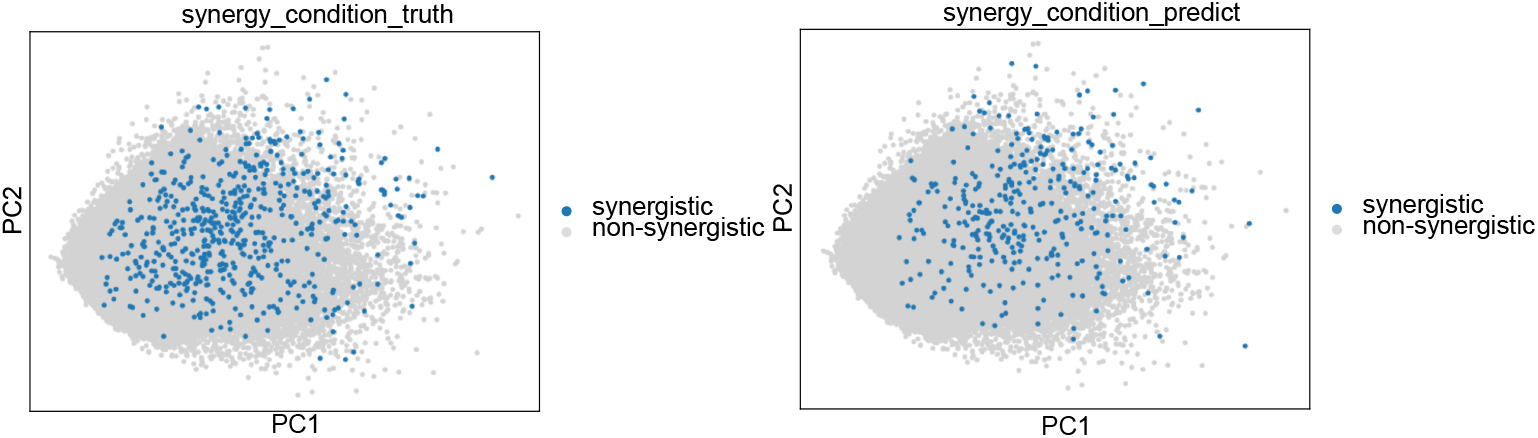
Visualization for the trained feature space of pre-training datasets (sub-sampled to 10 %). (a) The PCA plot from the outputs based on hidden layers of BAITSAO, colored by the ground truth synergistic information. (b) The PCA plot from the outputs based on hidden layers of BAITSAO, colored by the predicted synergistic information.

**Supplementary Fig. 10.**
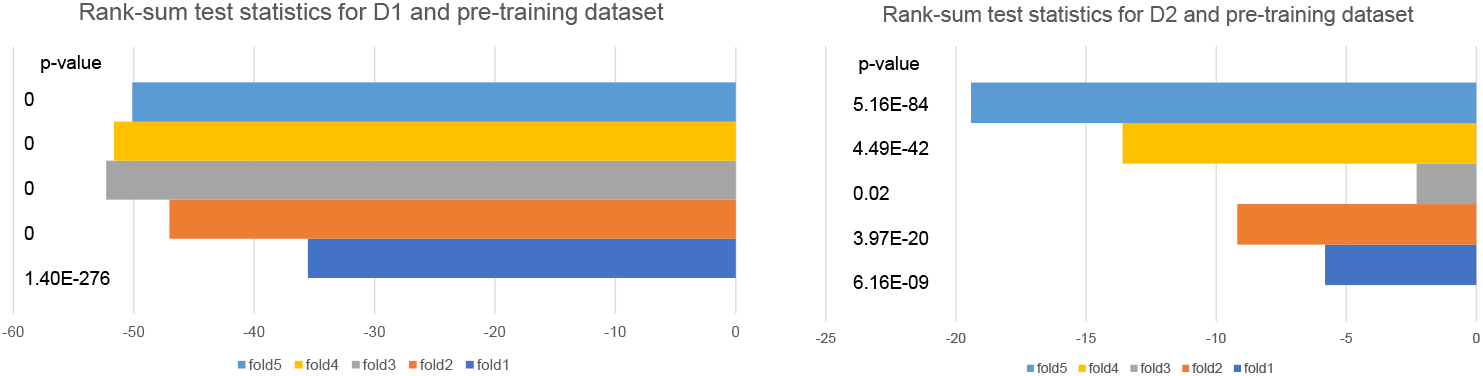
Statistics of the Rank-sum tests between the pre-training dataset and fine-tuning datasets. We included the two-side p-value for each comparisons. (a) The statistics of different folds by checking whether samples from D1 and the pre-training dataset come from the same distribution or not. (b) The statistics of different folds by checking whether samples from D2 and the pre-training dataset come from the same distribution or not.

**Supplementary Fig. 11.**
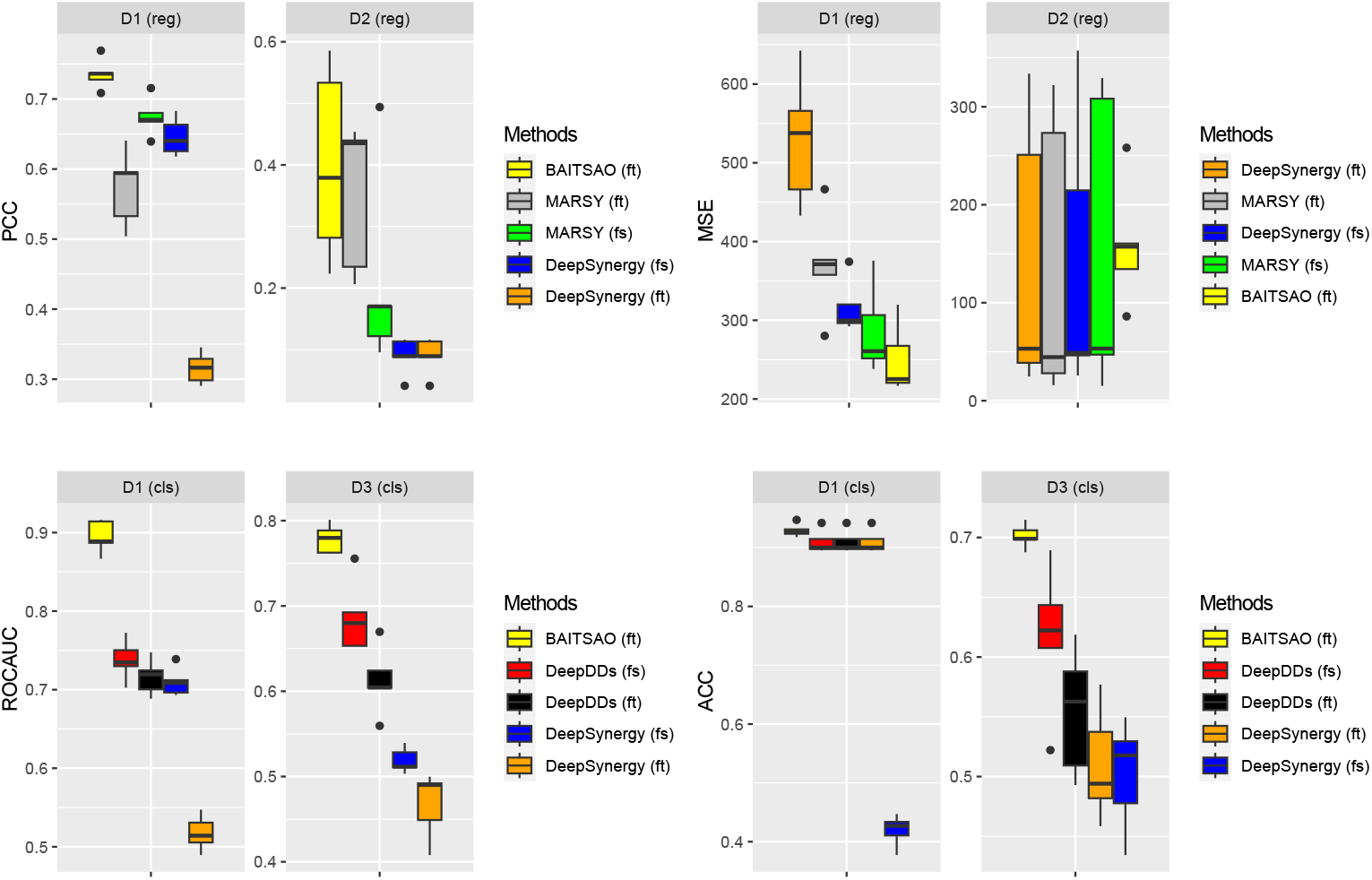
Comparisons of deep-learning-based models with pre-training (ending with ft) and from scratch (ending with fs) for two tasks across three different datasets. Data are presented in boxplots (n=5 per group; center line, median; box limits, upper and lower quartiles; whiskers, up to 1.5*×*interquartile range; points, outliers).

**Supplementary Fig. 12.**
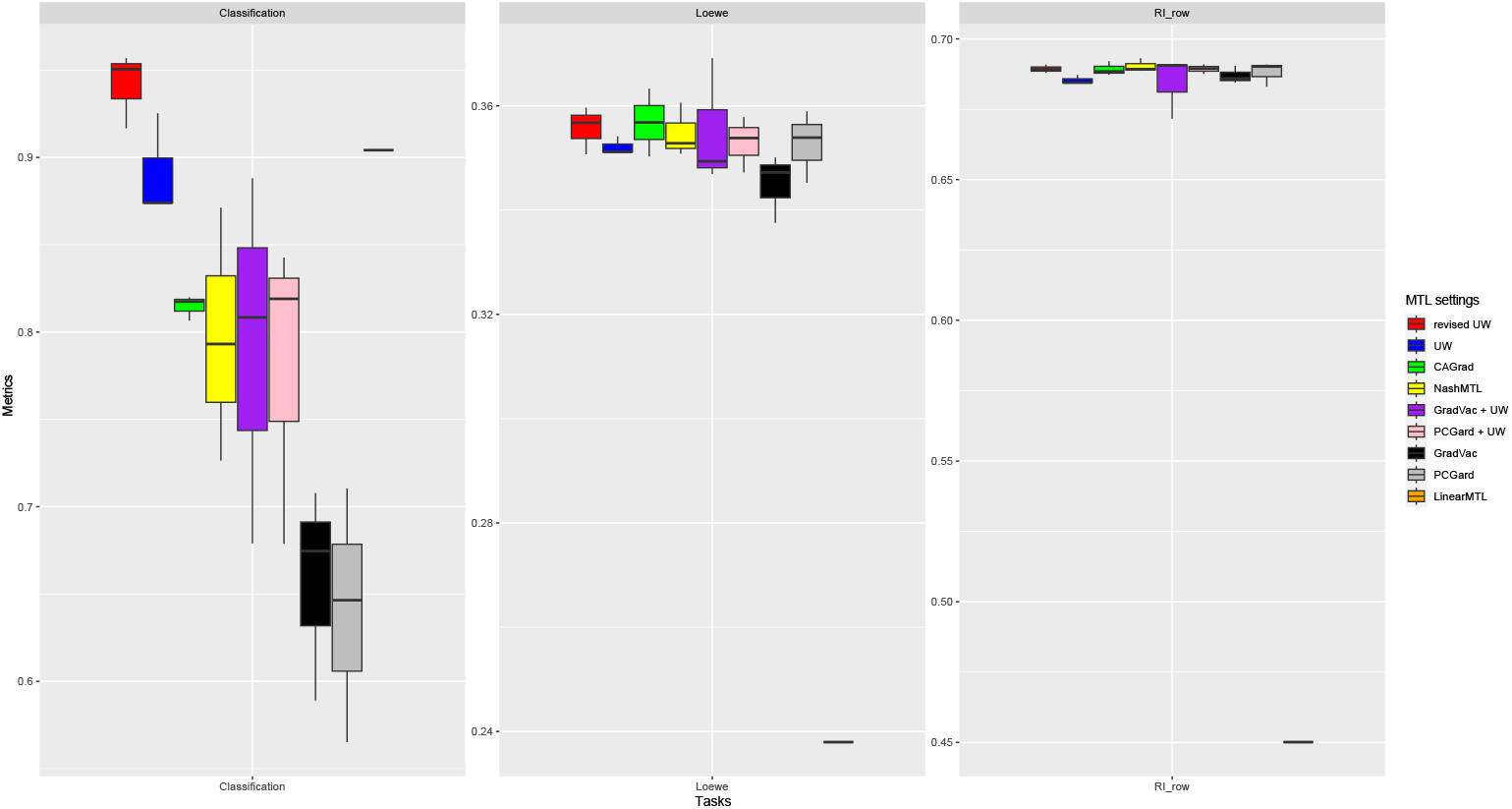
Ablation tests for MTL strategies. Here *PCGrad* +*UW* means we combine PCGrad with UW, and *GradVac*+*UW* means we combine GradVac with UW. *revised UW* represents the modified UW in this manuscript. For Loewe and RI row, we present scores of PCC. For classification, we present scores of ROCAUC. Data are presented in boxplots (n=3 per group; center line, median; box limits, upper and lower quartiles; whiskers, up to 1.5× interquartile range; points, outliers).

**Supplementary Fig. 13.**
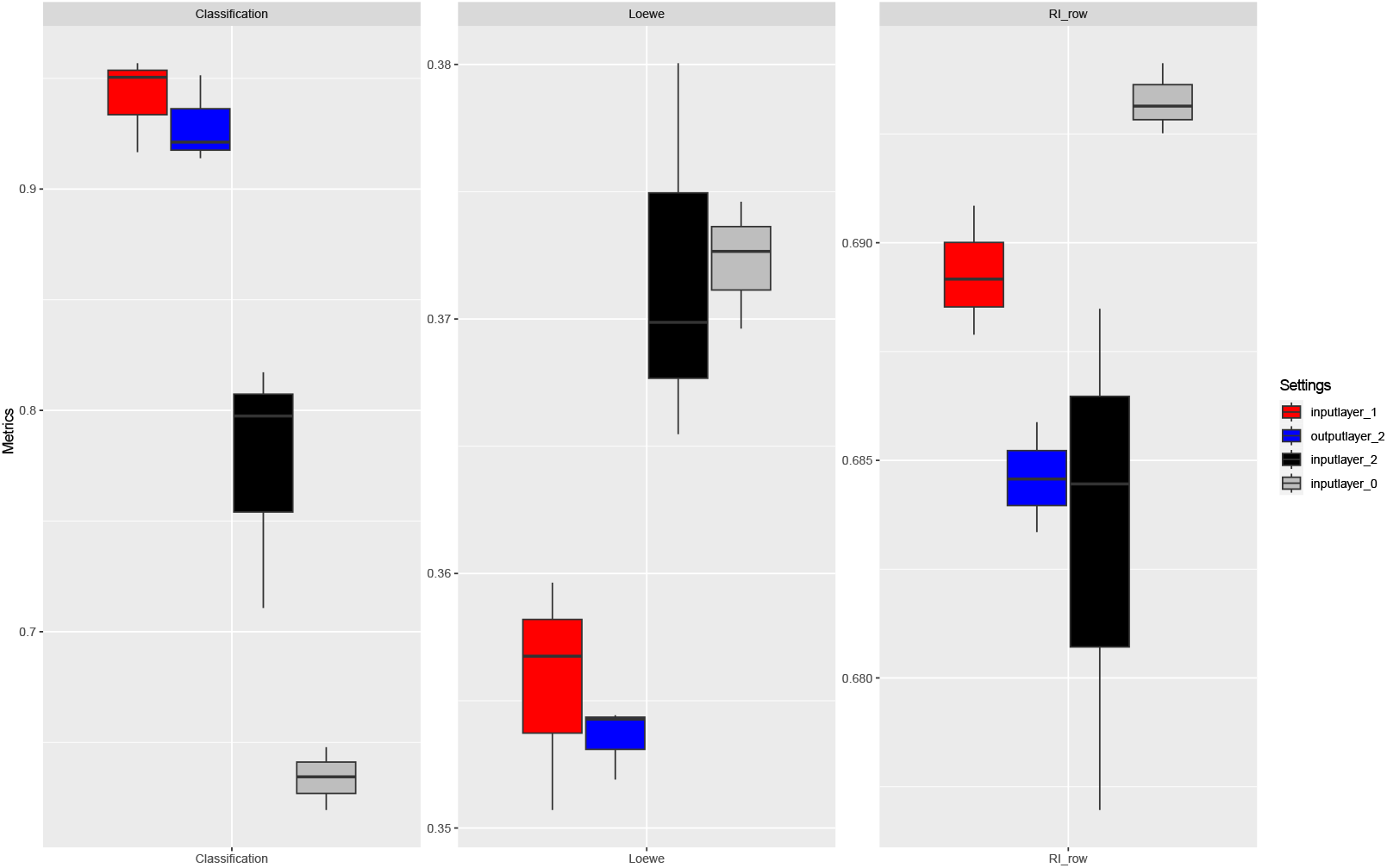
Ablation tests for the number of task-specific layers. Here *inputlayer 1* represents using one layer for processing input data and one layer for model’s output, and it is our final choice. *outputlayer 2* represents using one layer for processing input data and two layers for model’s output. *inputlayer 2* represents using two layers for processing input data and one layer for model’s output. *inputlayer 0* represents we did not set task-specific layers for input data. Data are presented in boxplots (n=3 per group; center line, median; box limits, upper and lower quartiles; whiskers, up to 1.5*×*interquartile range; points, outliers).

**Supplementary Fig. 14.**
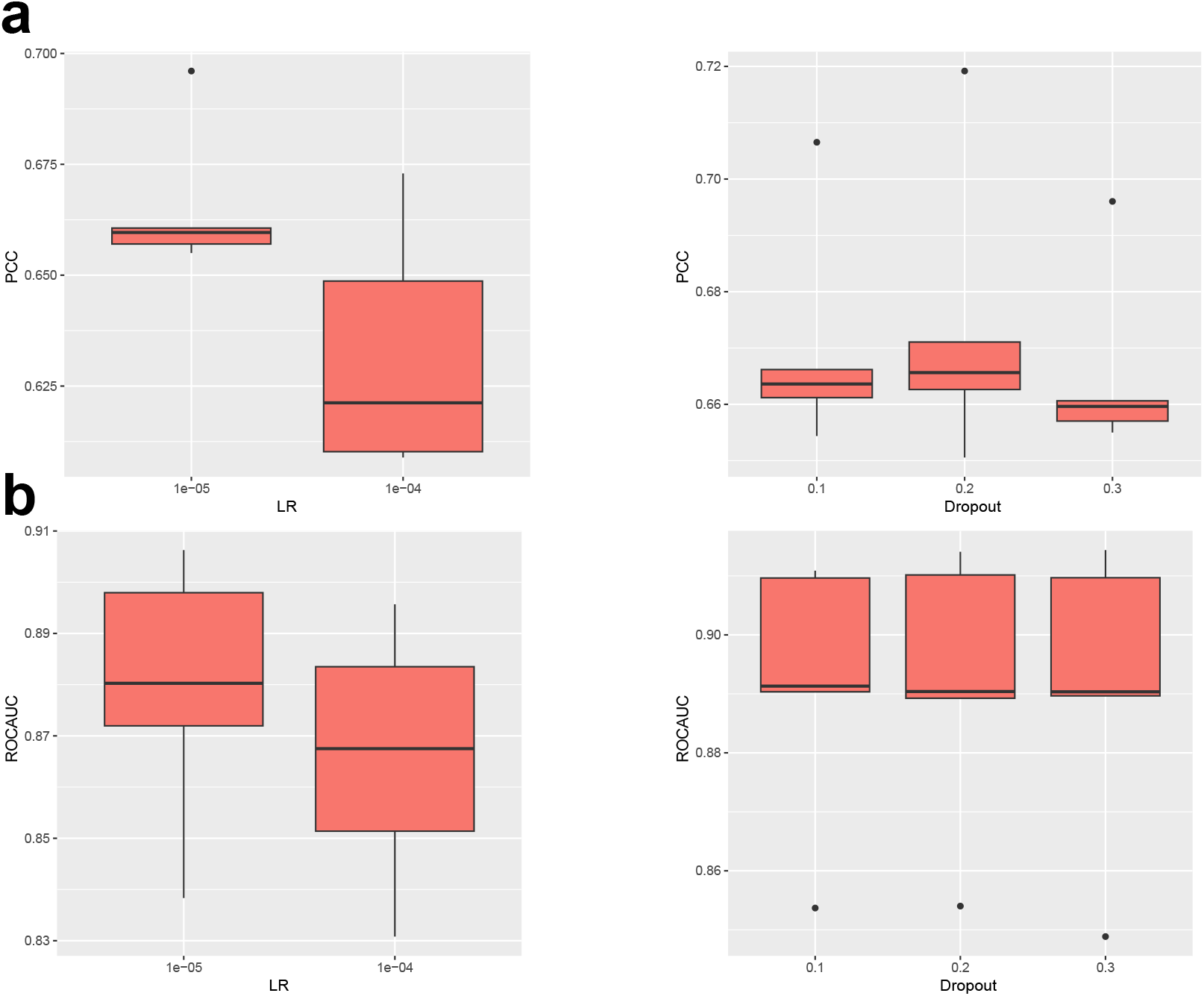
Results of hyper-parameter tuning of BAITSAO for D1. (a) The tuning results for the regression task. (b) The tuning results for the classification task. Data are presented in boxplots (n=5 per group; center line, median; box limits, upper and lower quartiles; whiskers, up to 1.5*×*interquartile range; points, outliers).

**Supplementary Fig. 15.**
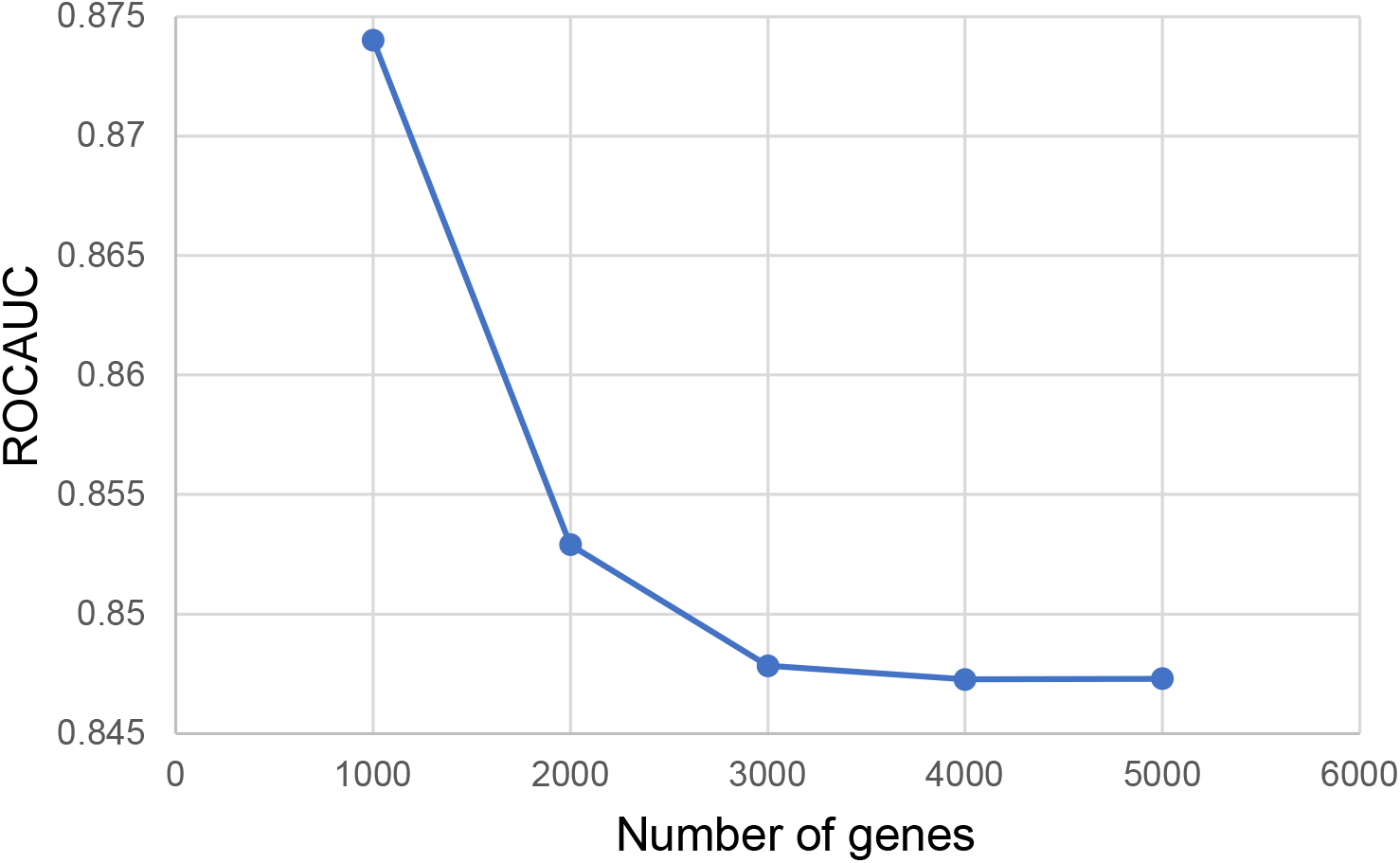
Results of tuning number of highly-variable genes for the analysis of explainability.

**Supplementary Fig. 16.**
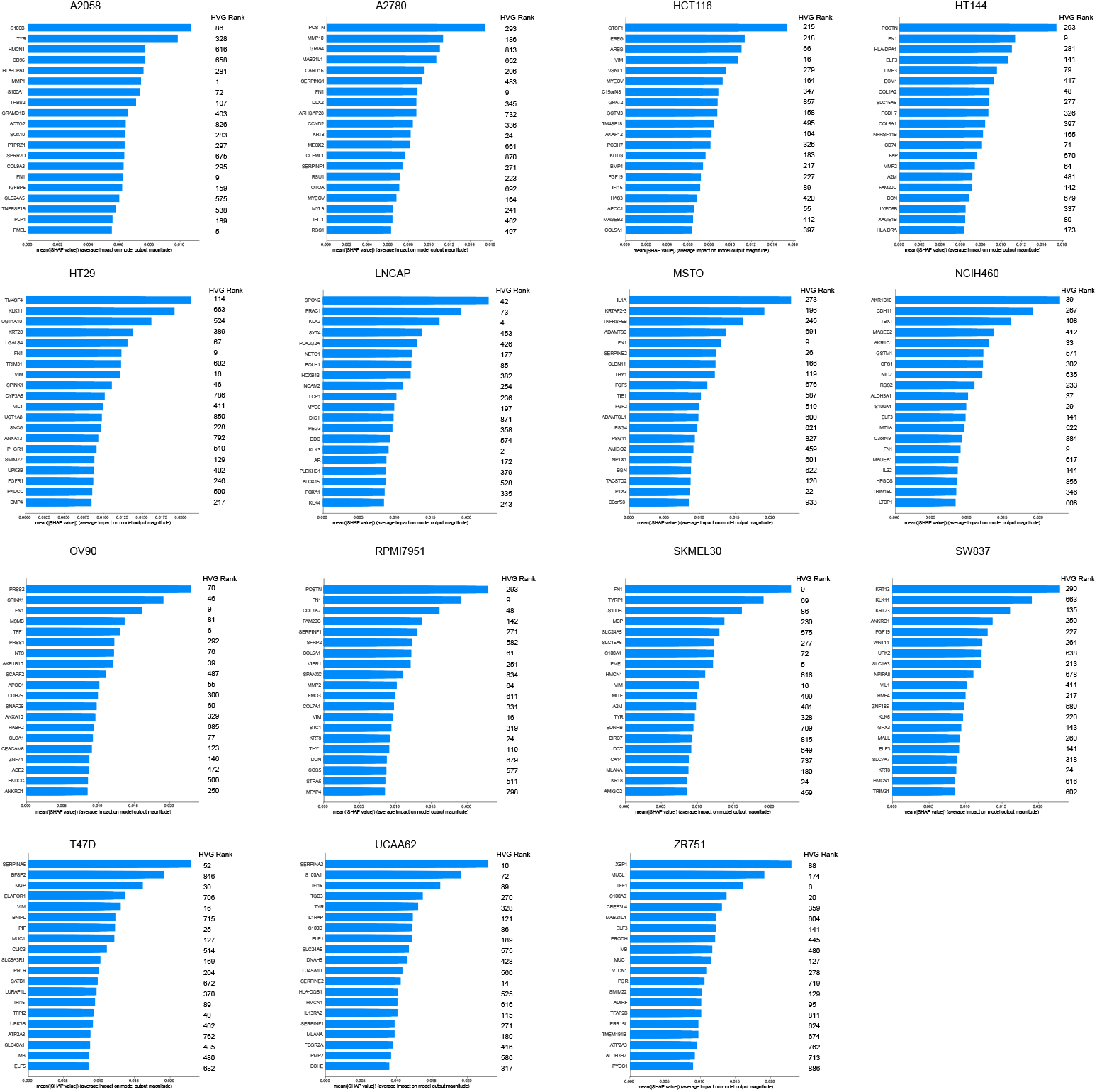
Visualization of important genes across different cell lines. We also annotate the rank based on variance for each gene.

**Supplementary Fig. 17.**
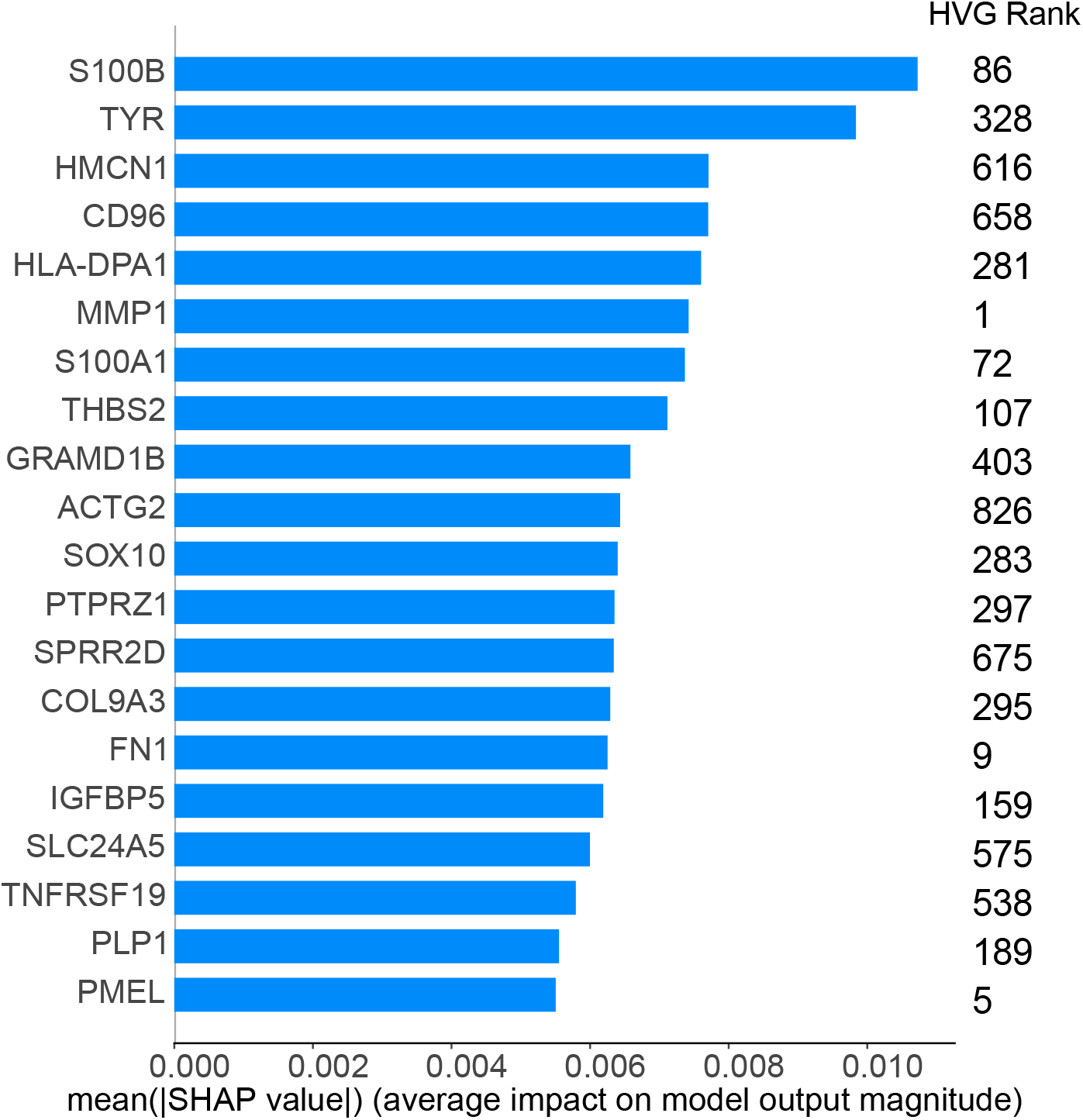
The explainability of BAITSAO for the combination: DEXAMETHASONE (drug)-DINACICLIB (drug)-A2058 (cell line). We also annotate the rank based on variance for each gene.

**Supplementary Fig. 18.**
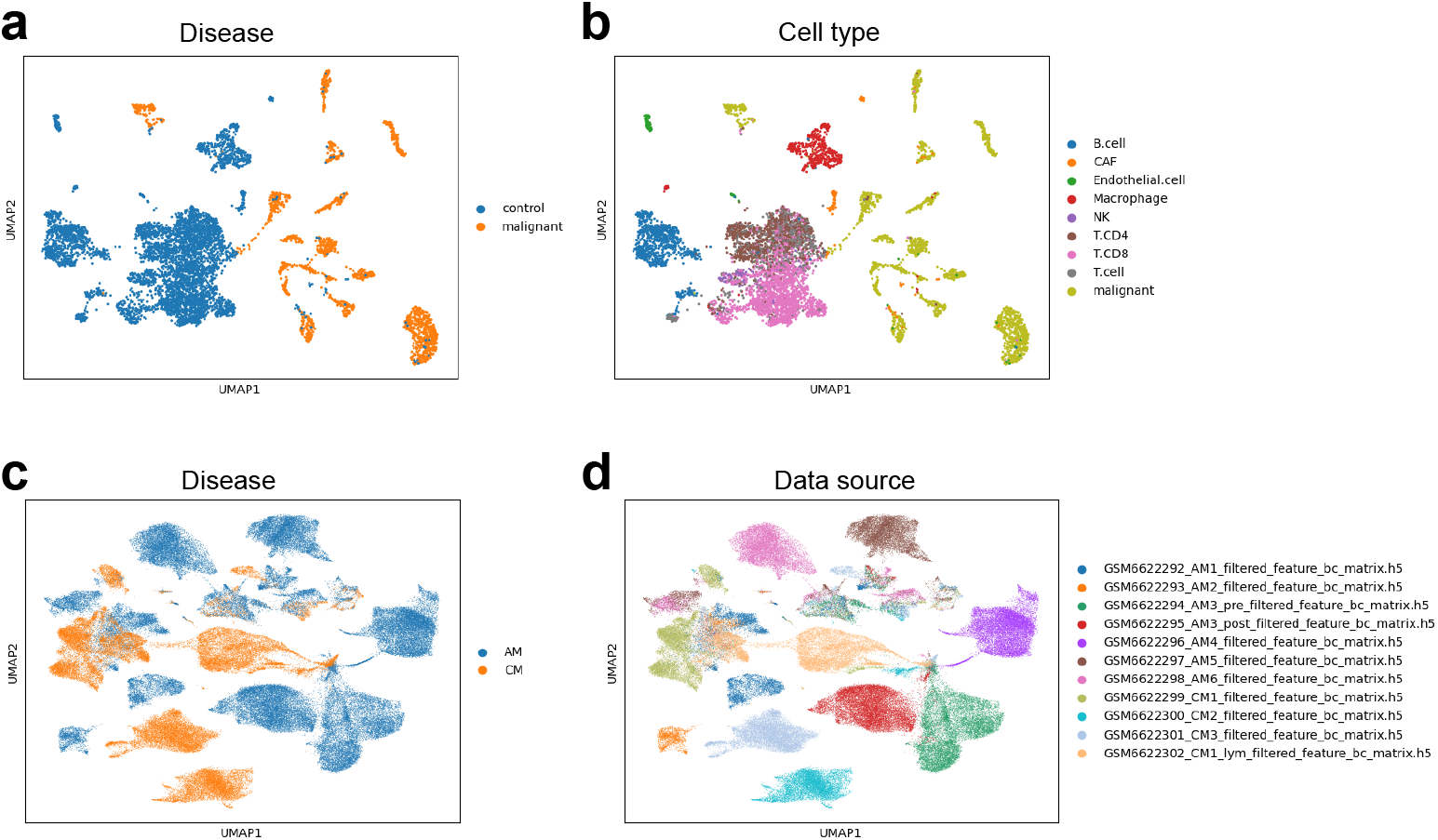
Visualization for two scRNA-seq melanoma datasets. (a) The UMAP plot for the scRNA-seq dataset colored by the conditions of cells from the diseased-control scRNA-seq dataset. (b) The UMAP plot for the scRNA-seq dataset colored by cell types from the diseased-control scRNA-seq dataset. (c) The UMAP plot for the scRNA-seq dataset colored by the conditions of cells from the AM-CM scRNA-seq dataset. (d) The UMAP plot for the scRNA-seq dataset colored by the sources of cells from the AM-CM scRNA-seq dataset.

## Footnotes

1 BAITSAO means collections of herbs and drugs in Chinese culture.

